# Conformational changes underlying electromechanical transduction in prestin resemble a transport transition in pendrin

**DOI:** 10.64898/2026.06.06.730624

**Authors:** Chenou Zhang, Richard Mariadasse, Jie Yang, Jun-Ping Bai, Joseph Santos-Sacchi, Dhasakumar S. Navaratnam, Oliver Beckstein

**Affiliations:** Center for Biological Physics, Arizona State University; Department of Physics, Arizona State University; Department of Neurology, Yale University School of Medicine; Department of Surgery (Otolaryngology), Yale University School of Medicine; Department of Neuroscience, Yale University School of Medicine; Department of Cell and Molecular Physiology, Yale University School of Medicine; The First Affiliated Hospital of USTC, Division of Life Sciences and Medicine, University of Science and Technology of China, Hefei, Anhui, 230001, P.R. China

## Abstract

Prestin (SLC26A5), a membrane protein in cochlear outer hair cells, drives electromechanical transduction essential for mammalian hearing. Unlike other SLC26 anion transporters, prestin functions as a voltage-dependent molecular motor, transitioning between contracted and expanded conformations. How this transition relates to the transporter cycle of SLC26 family members remains unclear. Here, multi-microsecond molecular dynamics (MD) simulations starting from a contracted conformation reveal a rapid, spontaneous transition to an expanded state that resembles the inward-facing conformation of the anion exchanger pendrin (SLC26A4 from mouse). An accompanying transmembrane area expansion is localized to the inner membrane leaflet. In line with this observation, reduced unitary sensor charge movement accompanies neutralization of charged residues localized near the inner leaflet. Simulations uncover a previously uncharacterized contracted conformation that resembles outward-facing pendrin; the MD also predicts an extracellular anion-binding site in prestin. A 3.27-Å resolution cryo-electron microscopy structure of prestin in the presence of thiocyanate anions confirms this site, which appears to be common in the SLC26 family, notably also in pendrin. Based on the simulations, the extracellular site in pendrin is involved in facilitating anion access to the canonical intracellular site in a conformation-dependent manner while in prestin, the intracellular site is not accessible from the extracellular side. Furthermore, like prestin, pendrin exhibits a non-linear capacitance, an indication of voltage-dependent conformational switching. Together, these findings suggest that prestin and pendrin share core structural and functional properties, notably parallels between expansion-contraction states and anion binding sites, though transition speeds and conformation-dependent ion-accessibility may differ.

## Introduction

Electromotility (eM) in the outer hair cells (OHC) of mammals is responsible for cochlear amplification, the principal mechanism underlying the exquisite sensitivity of mammalian hearing. Bill Brownell’s seminal observation of OHC eM ^1^ was followed by discoveries that the phenomenon was driven by 1) membrane voltage ^2^ and 2) expansion-contraction of the membrane surface area rather than from cytoskeletal elements ^3^. The conformational movements of a molecular motor protein (now known as prestin, see below) within the membrane result in a voltage-dependent change in membrane capacitance, known as non-linear capacitance (NLC). NLC results from voltage-sensor charge movements, being the first derivative of its charge-voltage (*Q*-*V*) relationship ^4,5^. Key parameters characterize NLC: *Q* the total charge moved during the protein’s reversible transitions from expanded to contracted state; *z* the unitary charge/valence of the motor; and *V*_ℎ_ the voltage at peak NLC, where half the motors are either contracted or expanded ^4,5^. Two models were proposed to explain prestin’s expansion and contraction: the area motor model ^6^ and the membrane bending model ^7^. The membrane bending model posits an expansion of the protein close to the inner leaflet of the membrane, whereas the area motor model is less specific and simply requires an increase in cross-sectional protein area. How these models relate to actual molecular structures of prestin in a realistic membrane environment remains an open question, but it is clear that prestin switches conformation between compact and expanded states in a voltage-dependent manner.

The cryo-EM structures of prestin (SLC26A5; PDB:7SUN ^8^, PDB:7LGU ^9^, and PDB:7S8X ^10^) resemble other members of the SLC26 anion transporter family. Prestin is a homodimer with each protomer possessing 14 transmembrane (TM) helices separated into a core domain with six TM helices, a gate domain with eight TM helices and a domain-swapped intracellular Sulphate Transporter and Anti-Sigma factor antagonist (STAS) domain (Figure 1a,b). Chloride anions modulate prestin electromechanics ^11–13^. For cryo-EM preparation in the presence of high Cl^−^ concentrations, prestin likely adopts a contracted state; that is, at these concentrations, *V*_ℎ_ of NLC is near −100 mV and thus at 0 mV (equivalent to an absent voltage gradient across either detergent micelles or nanodiscs), contracted conformations would predominate. The structures of prestin in Cl^−^ most closely match the intermediate structure of the related SLC26A9 (PDB:6RTF) in the transporter cycle ^14^. Members of the SLC26 family, like those of the related SLC6 and SLC4 transporter families, are believed to perform secondary active transport according to the alternating access mechanism ^15–17^. They undergo an elevator-like movement in their transporter cycle with the core domain moving relative to the gate domain; deduced conformations are (1) inward facing (IF), (2) intermediate (possibly occluded) and (3) outward facing (OF) ^18^. In the presence of Cl^−^, most members of the SLC26 family adopt an IF or intermediate structure ^8–10,14,19–24^ (Figure 1c). An internal anion binding site has been identified in all SLC26 family members at the interface of the core and gate domain and is accessible to the cytoplasm in the IF structures. Two groups also identified prestin structures that were interpreted to be in the expanded state in the presence of salicylate ^9,10^, since salicylate shifts *V*_ℎ_ in a depolarizing direction (in addition to reducing NLC while unexpectedly preserving a linear component of eM ^25^). In these structures, the internal Cl^−^ binding site was displaced along the membrane normal towards the cytosolic side, and both groups proposed that the expansion/contraction cycle in prestin replicated the transporter cycle of the other SLC26 family members, in line with the same hypothesis put forward earlier by Oliver et al. ^11^ and Schaechinger and Oliver ^26^ (although the original idea that Cl^−^ would have to be involved during the cycle as the voltage sensor ^11^ is now disproved ^9^). So far, one family member, pendrin (murine Slc26a4), has a OF structure in the presence of both its anion substrates, Cl^−^ and HCO^− 20^ (Figure 1d). In this structure, an external anion binding site was identified, which was located between the canonical intracellular anion binding site and the extracellular side at the interface of the core and gate domains and accessible to the extracellular environment ^20^. Thus, pendrin is the only SLC26 transporter with the two major conformations of the classical alternating access transport cycle resolved with physiological substrates and can therefore provide us with a structural framework in which to characterize the structurally homologous prestin.

**Figure 1.**
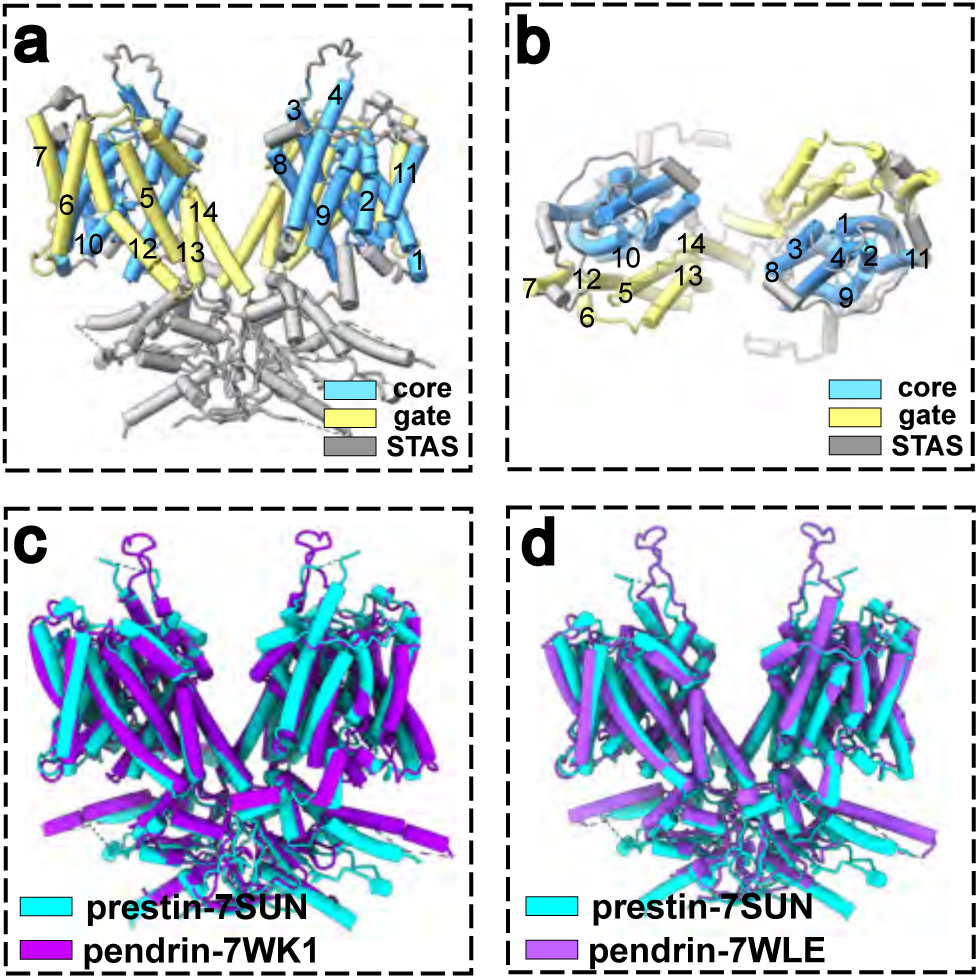
Architecture of prestin (SLC26A5) and comparison to pendrin (SLC26A4). **a** Prestin homodimer (PDB:7SUN) with core domain in blue and gate domain in yellow. Transmembrane helices (TM1-TM14) are labeled. The cytosolic STAS domain is colored gray. **b** Top view from the extracellular side of prestin. **c** Comparison of the structures of compact prestin (PDB:7SUN, cyan) and inward-facing pendrin (PDB:7WK1, magenta) after structural superposition on the gate domains. **d** Comparison of the structures of compact prestin (PDB:7SUN, cyan) and outward-facing pendrin (PDB:7WLE, magenta).

To better understand how prestin’s expansion/contraction occurs we performed molecular dynamics (MD) simulations using prestin from gerbil (*Meriones unguiculatus*) in a compact conformation (PDB:7SUN ^8^) as a starting point. Our simulations confirm that the expansion/contraction of the protein partially parallels the transporter cycle of pendrin and that the transition from the compact to the expanded state occurs rapidly, on the sub-microsecond to microsecond timescale. This intrinsic structural lability is compatible with prestin operating at very high, even ultrasonic, frequencies, in line with mechanical and electrical interrogations of prestin in OHCs ^27–33^, although our equilibrium simulations do not by themselves establish its voltage-driven operating bandwidth. Expansion due to the movement of the core domain is primarily restricted to the inner leaflet of the membrane. Furthermore, the simulations identify a new stable contracted conformation, compact-2, that resembles OF pendrin, and we additionally predict an extracellular anion binding site in prestin equivalent to the one in pendrin. We confirm this binding site through a new cryo-EM structure at 3.27 Å resolution in the presence of thiocyanate. Our simulations also indicate that pendrin expands its area when switching from the OF to the IF conformation and together with electrophysiologically measured NLC, our results suggest that prestin and pendrin share functional and structural features sufficient for electromechanical transduction.

## Results

Physiologically, prestin operates out of equilibrium: The OHC membrane potential changes continuously, and prestin responds by switching between at least two major macrostates that can be distinguished by their in-membrane cross-sectional area: a *contracted state* (C) and an *expanded state* (E), with depolarization favoring the contracted state and hyperpolarization the expanded state. The same two macrostates, however, are also accessible at thermal equilibrium. In the absence of a driving force, C and E interconvert spontaneously and the protein samples a Boltzmann equilibrium C ⇌ E whose populations are set by the free-energy difference between the states. External conditions—the membrane potential, but also Cl^-^ concentration, binding of small molecules, temperature, pressure or tension, and membrane lipid composition—shift this equilibrium, yet regardless of these conditions a finite probability exists to observe either conformation. For example, even though at a depolarizing potential (e.g., 0 mV) the equilibrium is shifted towards the contracted state under physiological conditions, a small fraction of prestin molecules will still occupy the expanded conformation. Consequently, even though all our simulations were performed under equilibrium conditions without any applied membrane voltage, we expect prestin to sample both macrostates, including the expanded state, and to undergo C ⇌ E transitions given sufficient sampling.

### In MD simulations prestin samples conformations resembling the IF and OF conformation of pendrin

Simulations starting from the experimentally determined compact structure of gerbil prestin (PDB:7SUN; Figure 2a) consistently showed relatively large conformational changes within tens to hundreds of nanoseconds as measured by root mean square deviation (RMSD) of the core domain after superposition on the gate domain (Figure 2c). We hypothesized that the conformational transition of prestin resembles part of the transition of pendrin for which IF and OF conformations have been experimentally determined ^20^. Thus, we constructed order parameters *ζ* and *φ* to quantify conformations of prestin relative to pendrin. After superimposing a protomer onto the gate domain of the pendrin IF reference structure (PDB:7WK1), the *ζ* order parameter measures the relative translational movement in the *z*-direction between the centers of mass of the core domains of prestin and pendrin, corresponding to the elevator movement along the membrane normal ^18^. The *φ* order parameter measures the rotational angle between the core domains of prestin and the IF pendrin reference structure, which in some elevator transporters governs access to the central binding site (Figure 2a, b).

**Figure 2.**
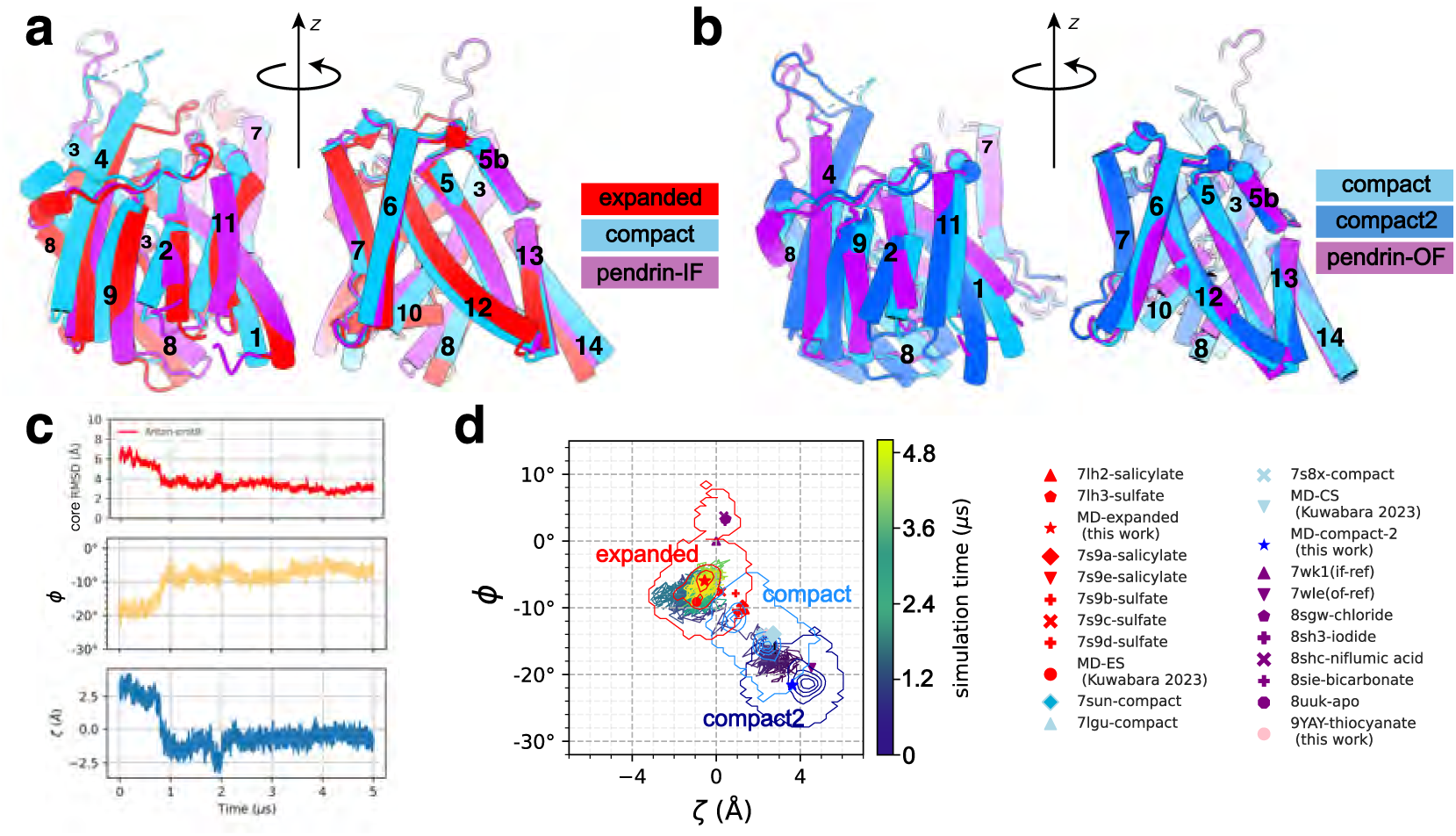
Prestin samples a wide range of conformations in MD simulations. **a** Inward-facing conformations: Structural superposition (on the gate domain) between pendrin (SLC26A4) in the IF conformation (purple, PDB:7WK1) and the expanded prestin conformation from MD (red); the intermediate compact prestin structure (light blue, PDB:7SUN) is provided for comparison. Left: View on the core domain (TM 1–4, 8–11). Right: View on the gate domain (TM 5–7, 12–14). **b** Outward-facing conformations: Structural superposition between pendrin in the OF conformation (purple, PDB:7WLE) and the compact-2 prestin conformation from MD (dark blue), with the intermediate compact conformation of prestin (light blue) also shown. Left: View on the core domain. Right: view on the gate domain. **c** Typical time series of structural order parameters (for protomer B in the 5-µs prestin-anton-0mV simulation) with pendrin in IF conformation as reference: Core-RMSD (RMSD of C*_α_* atoms in the core-domain after superposition on the gate domain), *φ* (rotation of the core domain towards the core domain of the reference structure) and *ζ* (elevator movement of the core domain towards the reference structure). **d** Projection of the same trajectory onto the 2D order-parameter space at 2.4-ns intervals and colored from dark to light according to simulation time from 0 to 5 µs. Contours corresponding to the expanded (red), compact (light blue), and compact-2 (dark blue) states that were obtained from a clustering analysis are shown for all simulation data. Experimental and simulation structures are marked in the 2D order-parameter space and colored by state (expanded in red; compact in light blue; compact-2 in dark blue; pendrin in magenta; new cryo-EM structure of gerbil prestin in the presence of thiocyanate in pink). The origin at *ζ* = 0 and *φ* = 0 is defined by the inward-facing pendrin reference structure.

All prestin simulations were initiated from the compact prestin structure (PDB:7SUN), which appears as the light-blue diamond in the order parameter plot (Figure 2d). Additional experimental compact-state structures (light-blue symbols) cluster tightly around the gerbil prestin structure, forming a well-defined compact region in the two-dimensional order-parameter space. When our MD trajectories were projected into the *ζ*, *φ* space, many of them showed clear transitions towards either IF or OF pendrin. For example, protomer B of prestin in the 5-µs long Anton-0mV trajectory (Table 1) transitions from the compact state toward the IF conformation of pendrin after 799.7 ns (Figure 2d, Supplementary Information Table S1), with *ζ* decreasing by ∼3 Å and *φ* increasing by ∼8^◦^.

**Table 1.** All-atom, explicit MD simulations of prestin and pendrin performed under equilibrium conditions Simulations are referenced in the text by their *simulation ID*. *PDB* indicates PDB ID of the experimental structure that the simulation is based on.

| simulation ID | protein | organism | PDB | atoms | time (μs) | MD package |
| --- | --- | --- | --- | --- | --- | --- |
| prestin-Anton-0mV<sup>1</sup> | prestin | gerbil | 7SUN<sup>2</sup> | 208096 | 5 | Anton 2 |
| prestin-Anton-repeat-1 | prestin | gerbil | 7SUN<sup>2</sup> | 208096 | 2 | Anton 2 |
| prestin-Anton-repeat-2 | prestin | gerbil | 7SUN<sup>2</sup> | 208096 | 2 | Anton 2 |
| prestin-Anton-repeat-3 | prestin | gerbil | 7SUN<sup>2</sup> | 208096 | 2 | Anton 2 |
| prestin-7lgu | prestin | human | 7LGU<sup>3</sup> | 288584 | 2 | Anton 2 |
| prestin-gmx-1 | prestin | gerbil | 7SUN<sup>2</sup> | 208096 | 2 | GROMACS 2021.3 |
| prestin-gmx-2 | prestin | gerbil | 7SUN<sup>2</sup> | 208096 | 2 | GROMACS 2021.3 |
| prestin-gmx-3 | prestin | gerbil | 7SUN<sup>2</sup> | 208096 | 2 | GROMACS 2021.3 |
| prestin-compact2-1<sup>5</sup> | prestin | gerbil | 7SUN<sup>2</sup> | 208096 | 1.1 | GROMACS 2021.3 |
| prestin-compact2-2<sup>5</sup> | prestin | gerbil | 7SUN<sup>2</sup> | 208096 | 1.2 | GROMACS 2021.3 |
| prestin-compact2-3<sup>5</sup> | prestin | gerbil | 7SUN<sup>2</sup> | 208096 | 1.9 | GROMACS 2021.3 |
| pendrin-inward | pendrin | mouse | 7WK1<sup>4</sup> | 244786 | 3 | GROMACS 2021.3 |
| pendrin-inward-repeat1 | pendrin | mouse | 7WK1<sup>4</sup> | 244786 | 1 | GROMACS 2021.3 |
| pendrin-inward-repeat2 | pendrin | mouse | 7WK1<sup>4</sup> | 244786 | 1 | GROMACS 2021.3 |
| pendrin-inward-repeat3 | pendrin | mouse | 7WK1<sup>4</sup> | 244786 | 1 | GROMACS 2021.3 |
| pendrin-outward | pendrin | mouse | 7WLE<sup>4</sup> | 247744 | 3 | GROMACS 2021.3 |
| pendrin-outward-repeat1 | pendrin | mouse | 7WLE<sup>4</sup> | 247744 | 1 | GROMACS 2021.3 |
| pendrin-outward-repeat2 | pendrin | mouse | 7WLE<sup>4</sup> | 247744 | 1 | GROMACS 2021.3 |
| pendrin-outward-repeat3 | pendrin | mouse | 7WLE<sup>4</sup> | 247744 | 1 | GROMACS 2021.3 |

<sup>1</sup> simulation from Bai et al.<sup>33</sup>
<sup>2</sup> Butan et al.<sup>8</sup>
<sup>3</sup> Ge et al.<sup>9</sup>
<sup>4</sup> Liu et al.<sup>20</sup>
<sup>5</sup> simulations were initialized from a trajectory snapshot from prestin-gmx-1 that contained the compact-2 conformation

Notably, during the first 500 ns, the protein samples conformations adjacent to OF pendrin before then moving rapidly towards IF pendrin. A number of experimental structures of prestin have been solved in the presence of ligands such as sulfate and salicylate that may shift the structural equilibrium towards the expanded conformation. Many of these structures are located in the order parameter plot towards IF pendrin (Figure 2d) but our MD simulations typically move beyond these structures. Kuwabara et al. ^34^ performed MD simulations of human prestin and observed a conformation that they classified as expanded; this conformation displays a larger *ζ* -shift but a smaller rotation angle *φ* than some of the experimental structures. Our simulations of prestin show a larger core-domain shift and larger core rotation than all the experimental putative expanded structures and appear to be closer to IF pendrin than any of the existing structures or models of expanded prestin.

We clustered all MD simulations for each protomer in the dimer separately based on structural similarity (see Methods and Supplementary Information Figure S1) and defined three major conformational clusters that we labeled *compact* (around the experimentally determined compact structure; PDB:7SUN), *expanded* (beyond the putative expanded structures), and *compact-2* (towards pendrin OF). Compact and compact-2 together constitute the contracted macrostate C, whereas the expanded cluster constitutes E. These clusters cover large (and partially overlapping) regions in the order parameter space (Figure 2d). This state classification approach was performed independently of the order parameter analysis and so these two approaches provide complementary views on the data. Although not all trajectory frames could be neatly classified (see Supplementary Information Figure S2), the classification allows us to increase the signal-to-noise level and focus on the large-scale conformational changes that are evident in the data.

Structural comparisons to the pendrin experimental structures support this classification (see Supplementary Information for a detailed comparison, including Supplementary Information Table S2 and S3). The expanded model (Figure 2a) is closer to IF pendrin (PDB:7WK1) than to the compact prestin structure (PDB:7SUN), with core-domain RMSDs after superposition on the gate domain of 2.8 Å and 5.1 Å, respectively, whereas the compact-2 model (Figure 2b) lies intermediate between compact prestin (4.5 Å) and OF pendrin (PDB:7WLE; 6.5 Å) (Supplementary Information Table S3). Because the gate domain superimposes closely in all comparisons (RMSD ≤ 1.9 Å, Supplementary Information Table S3), these differences reside almost entirely in the position and orientation of the core domain, which in IF pendrin is shifted and rotated somewhat further than in our expanded model.

### Prestin (and pendrin) expand at the inner leaflet of the membrane

We developed a new method based on a stratified 2D Voronoi tessellation to quantify the cross-sectional area of a protein in the membrane from MD simulations (see Methods) to assess the functional relevance of the conformations in the different states (Figure 3). The average cross-sectional area profile *A*(*z*) is larger in the *expanded* state than in the *compact* one (Figure 3b). The largest difference (on the order of 150 Å^2^) is located at the inner leaflet, marked by a high density of lipid acyl chains. Differences in the headgroup region are less pronounced and of lower relevance as judged from the larger statistical error. Thus, the simulations indicate clearly that prestin’s expansion is largely restricted to the TM regions closest to the inner leaflet of the membrane (Figure 3b) and is primarily due to the movements of TM helices 4, 9, 2, and 11 and the outward movement of TM 1 (Supplementary Information Figure S3). Consistent with the simulation data, as previously shown ^8,9^, the location of charged residues that reduced *z*, the unitary moving charge of prestin (see equation (2)), when neutralized were found to localize closer to the inner leaflet of the membrane in the TM segments of the protein (Figure 3d, e). We expanded on these data and show that several additional charged residues in TM1 and 6 that lie closer to the inner leaflet also reduced *z* when neutralized (Supplementary Information Table S4). Thus, to date 15 mutated charged residues were found to reduce *z*, of which 14 residues lie in proximity to the inner leaflet (Figure 3d,e and Supplementary Information Table S4). In contrast, 20 mutated charged residues were found to not affect *z* and are approximately equally distributed at the inner and outer leaflets with 12 lying in proximity to the outer leaflet (Supplementary Information Table S4).

**Figure 3.**
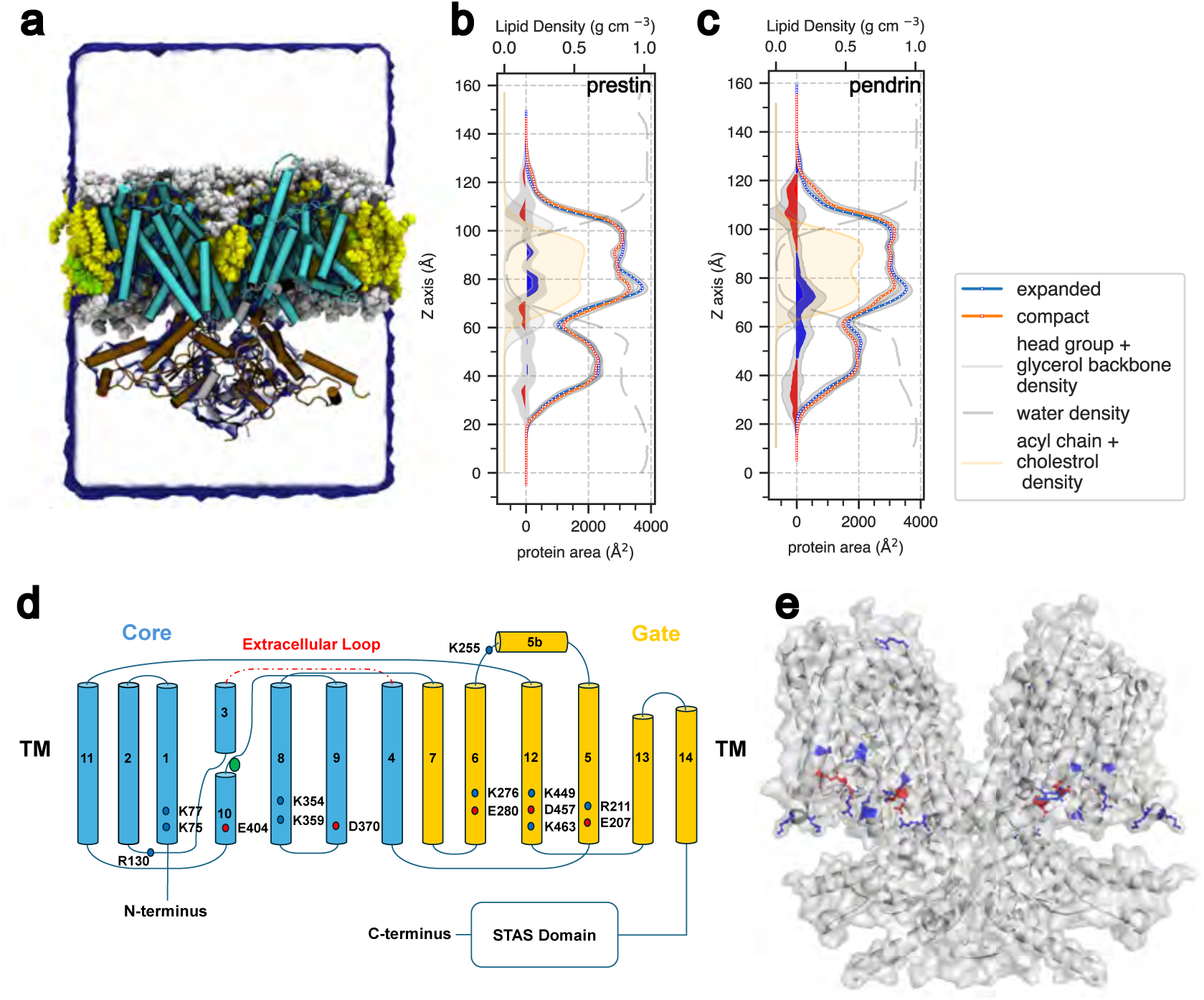
Protein area difference in MD simulations. **a** MD simulation system of prestin (PDB:7SUN), with the transmembrane domain shown in cyan, the STAS domain in brown, surrounding POPC lipids in gray (head groups and glycerol backbone) and yellow (acyl chains), and cholesterol (green). A contour of the water density (blue) outlines the simulation box. **b** Prestin area difference. Averaged area profiles along the *z*-axis from the prestin-Anton-0mV simulation, with compact (red) and expanded (blue) states computed separately; standard deviations for each state are shown in gray. The difference between the compact and expanded area profiles is plotted on the left. Lipid and water densities are shown by group: water (gray dashed), lipid headgroup+backbone (gray), and acyl chains or cholesterol (yellow). **c** Pendrin area difference. **d** General topology of prestin. Mutated residues that affect NLC are labeled and colored (basic in red, acidic in blue). **e** The location of these mutations in prestin (see Supplementary Information Table S4 for the list of residues).

The maximum cross-sectional area of the dimer varies across simulations from about 3100 Å^2^ to slightly less than 4100 Å^2^. When the area distributions are separated by the number of protomers classified as expanded, a clear linear trend emerges (Figure 4c). We used a simple linear model

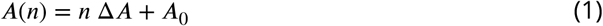

to relate the conformational state of a protomer (obtained from the structural clustering) to the functionally relevant maximum cross-sectional area (see more details in Methods). In equation (1), *n* is the number of protomers in the expanded conformation, Δ*A* the area expansion per protomer, and *A*_0_ the dimer area in the fully contracted *n* = 0 conformation. From a least-squares fit of all classified trajectory frames to the model (Figure 4c) we find that on average each protomer expands by Δ*A* = 149.6 ± 0.3 Å^2^ when undergoing the transition from C (contracted: compact or compact-2) to E (expanded); the average area of the fully contracted dimer is *A*_0_ = 3424.0 ± 0.3 Å^2^ (errors are standard errors of the parameter estimate from the least-squares fit). The relative expansion is 2Δ*A*∕*A*_0_ = 8.8%. The area for each protomer does not differ between the compact and compact-2 states (Figure 4d) but both differ from the expanded state, thus indicating self-consistency of our model in which we combined compact and compact-2 states into a single contracted C state.

**Figure 4.**
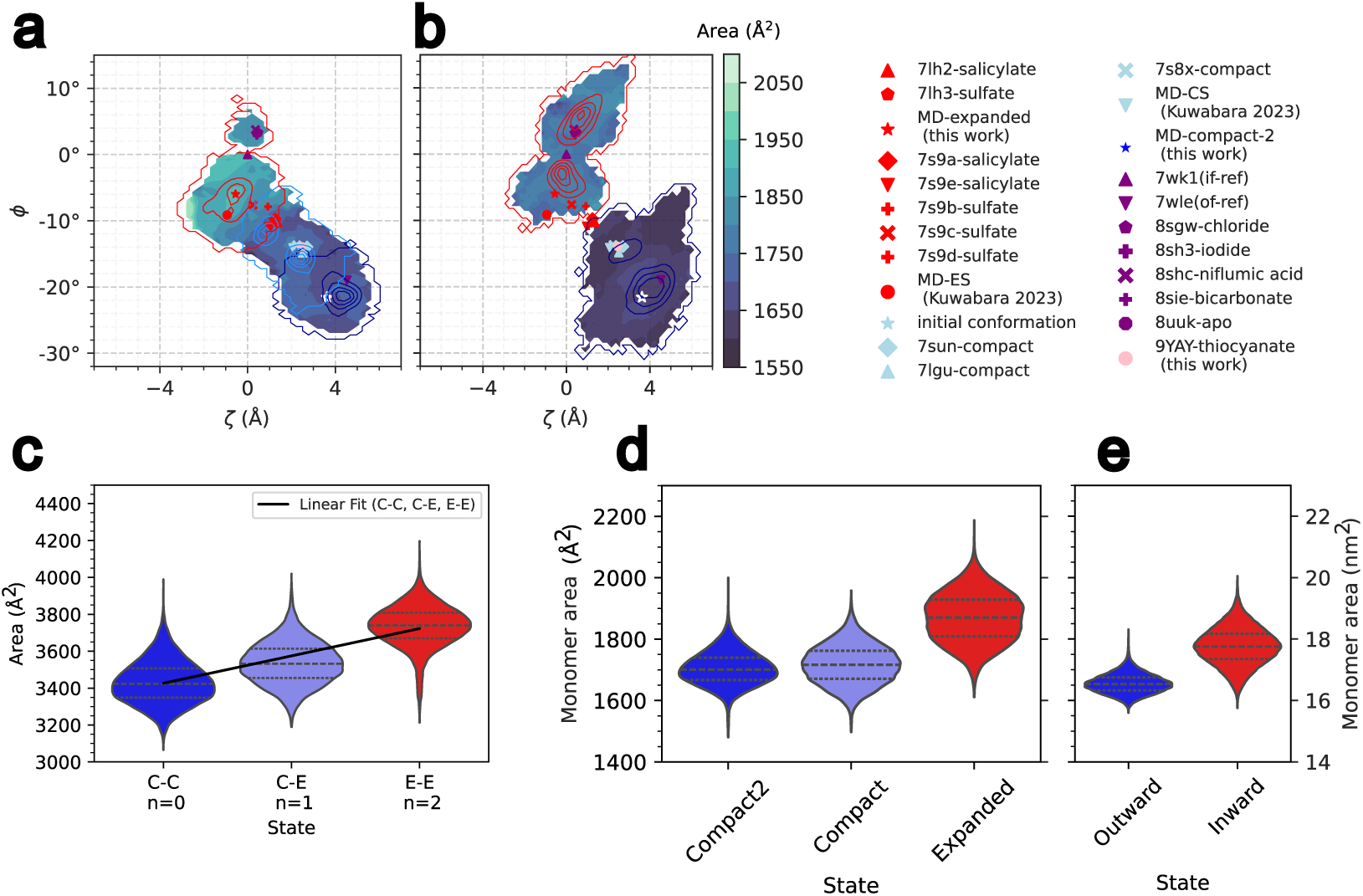
Area of prestin and pendrin depends on the conformation. **a** Projection of the protomer area of prestin in MD simulations on the *ζ*, *φ* 2D order-parameter space. Known experimental structures and contours for classified expanded (red), compact (light blue) and compact-2 (dark blue) conformations are plotted as in Figure 2d. **b** Projection of the protomer area of pendrin from MD simulations. **c** Violin plot of dimer-area distributions from all prestin MD simulations, separated by dimer-state combinations: expanded/expanded (EE), expanded/contracted (EC), and contracted/contracted (CC; contracted includes compact and compact-2). A linear fit is shown for the relationship between the number of expanded protomers and the resulting dimer area. **d** Violin plot of prestin monomer maximum area distributions separated by conformational state. The mean areas for each state are: compact-2 1704 Å^2^, compact 1717 Å^2^, expanded 1869 Å^2^. **e** Violin plot of pendrin monomer maximum area distributions separated by conformational state.

The protomer area, projected onto the *ζ*, *φ* order parameters, shows that conformations more similar to OF pendrin (compact-2) or to compact prestin structures (compact) have smaller cross-sectional areas than conformations near IF pendrin (expanded) (Figure 4a). The area increases more steeply in the region where compact and expanded states overlap and where many of the experimental structures are located that have been described as “expanded”. Our analysis directly links the large-scale domain movement of the core domain to the functionally relevant cross-sectional area changes.

For comparison, we also performed MD simulations of pendrin in IF and OF conformations (Table 1). Over long 3-µs simulations and multiple shorter ones (3 × 1 µs) starting in either conformation, pendrin did not undergo any large conformational changes, as judged by RMSD and order parameters (Figure 4b). The cross-sectional areas of the TM regions of pendrin also show expansion closer to the inner leaflet of the membrane in the IF structure compared to the OF structure (Figure 3c). With the same linear model, the area increase from OF to IF per protomer is Δ*A* = 116.7 ± 0.2 Å^2^; the average area of the fully OF dimer is *A*_0_ = 3313.7 ± 0.3 Å^2^, which is comparable to what we observed for prestin and thus we may identify the pendrin IF conformation with an expanded state and the OF conformation with a contracted state (Figure 4e).

### Prestin shows rapid conformational transitions

For prestin (but not pendrin), we observed that each protomer could change its conformation rapidly when the MD simulations were initialized from the equilibrated compact structure. For example, protomer A in the Anton-0mV simulation had entered the transition region between compact and expanded states near the conformations of the sulfate and salicylate-stabilized expanded-like structure within less than a nanosecond (Supplementary Information Figure S4a) while protomer B had moved slightly towards the compact-2 region. Given that each protomer in almost all simulations eventually transitioned to the expanded state, we recorded the first-passage time for the first arrival in the expanded state (Supplementary Information Table S1). Because no reverse transition from expanded to either contracted state was observed in any simulation, these data characterize only the compact → expanded transition of the full equilibrium interconversion. Treating the first-passage times as a single-exponential process to estimate the transition rate with Bayesian inference ^35^ yields an order-of-magnitude rate of 2.2 µs^−1^ (95% confidence interval 1.3 to 3.8 µs^−1^; Supplementary Information Figure S5), corresponding to a characteristic timescale of about 0.5 µs (0.3 to 0.8 µs confidence interval). Given the large variability between protomers and trajectories (Supplementary Information Table S1) and the absence of reverse transitions, this value should be interpreted as an approximate timescale for the compact-to-expanded relaxation under our equilibrium conditions, not as a voltage-driven switching frequency. It nonetheless indicates that the compact-to-expanded structural change is intrinsically fast, occurring on the submicrosecond to microsecond timescale.

To test if these fast transitions were specific to our simulations starting from the compact gerbil structure (PDB:7SUN), we also performed a 2-µs simulation starting from the highest-resolution compact structure of human prestin (2.30 Å; PDB:7LGU) ^9^ (Table 1). In these simulations both protomers transitioned rapidly from the compact conformation towards the expanded conformation (Supplementary Information Figure S6), with protomer B entering the expanded state after less than 40 ns of the start of the simulation and protomer A after 253.7 ns (Supplementary Information Table S1). Thus, our control simulation with human prestin confirmed that the rapid transitions towards the expanded conformation were not species-specific and unlikely due to the resolution of the starting structure.

### Pendrin presents NLC

Since prestin’s movement to the IF expanded form closely resembled the IF structure of pendrin and since previous data have shown pendrin to have measurable NLC ^36^, we sought to confirm the presence of NLC. Our intent was to measure NLC of pendrin as a confirmatory measure of its voltage-driven motor function. We found that pendrin exhibited a hyperpolarizing shift in NLC *V*_ℎ_ in chloride solution, compared to prestin. *V*_ℎ_ (−240 ± 23 mV) was inferred using a two-state Boltzmann model (Figure 5a, b). We could not directly observe the voltage at the peak in capacitance since it is beyond the measurement limits of the amplifier. To further validate this finding, we repeated the recordings in sulfate solutions, since sulfate causes a depolarizing shift in prestin’s *V*_ℎ_, reasoning that pendrin would show a similar depolarizing shift with 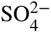. As shown in Figures 5a and 5c, the NLC peak in sulfate solution is indeed shifted, and estimated to be −217 ± 10 mV. These data complement prior results ^36^ that pendrin shows NLC, but more hyperpolarized than prestin.

**Figure 5.**
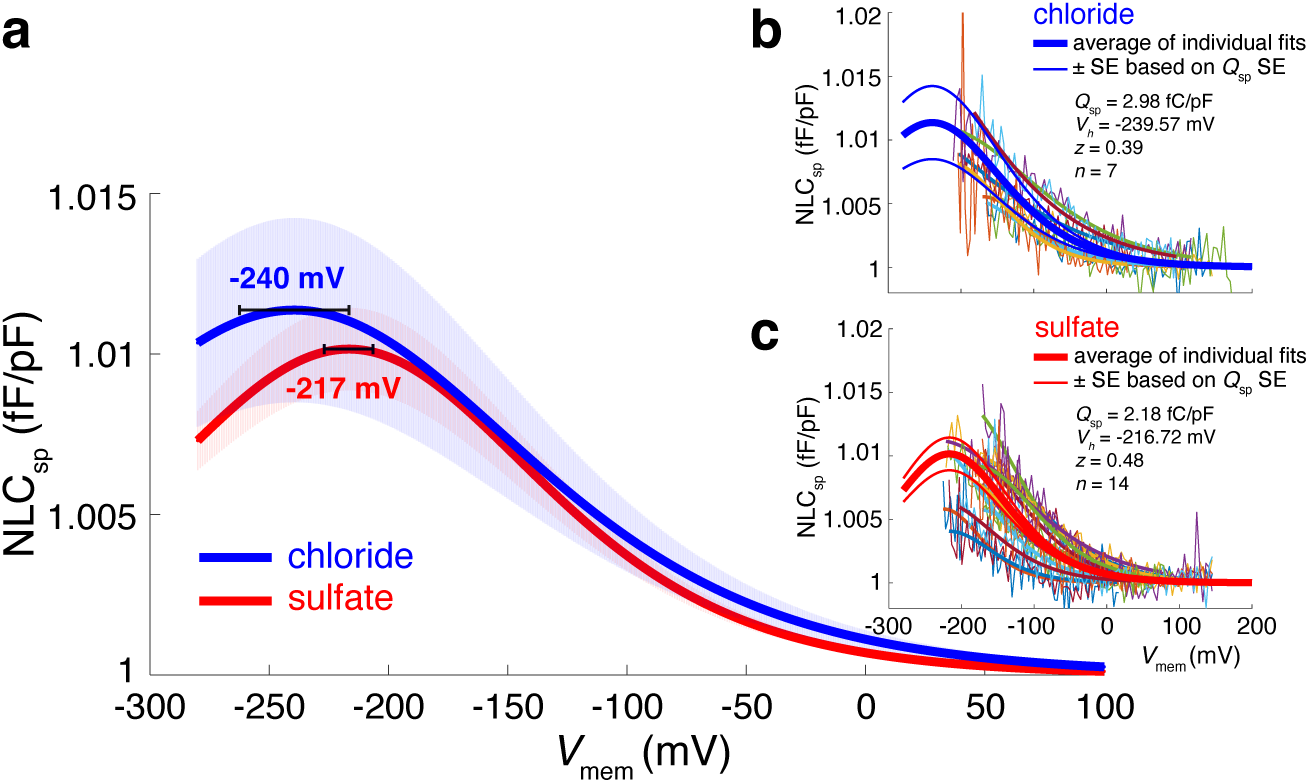
NLC measurement of pendrin in chloride (blue) and sulfate (red). **a** Averaged Boltzmann fits. Error bands indicate the standard error of the averaged curves shown in b and c. **b** Individual NLC measurements in chloride solution (n=7). **c** Individual measurements in sulfate solution (n=14). Different colors represent individual recordings. A two-state Boltzmann model was used to fit the data. Averaged NLC parameters are indicated in b and c.

### MD simulations identify an extracellular anion binding site in prestin

In previous work we established that Cl^−^ ions can freely access the canonical intracellular anion binding site of prestin (IC site), especially once the transition towards the expanded state has occurred ^33^, thus establishing the expanded conformation as an IF open state. In the MD simulations, we observed contracted states with and without Cl^−^ ions bound to the IC site. Exchange of Cl^−^ ions between the IC site and the intracellular compartment appears to be somewhat more frequent in the compact state than in the compact-2 state, but neither state appears to be fully occluded to Cl^−^ ions. After closer inspection, the simulations also revealed frequent binding of Cl^−^ ions to a site connected to the extracellular compartment around residues Lys227 and Arg236 (Figure 6c) with clear chloride ion density (Supplementary Information Figure S7c). The primary residues interacting with the anion in the EC site are Arg236 with average coordination number 0.45 ± 0.21, Lys227 (0.54 ± 0.18), Lys235 (0.28 ± 0.13), and Ser224 (0.25 ± 0.19; see Supplementary Information Table S5). Binding was observed independently of the conformation of the protomer and therefore the coordination numbers are averaged over all simulations.

**Figure 6.**
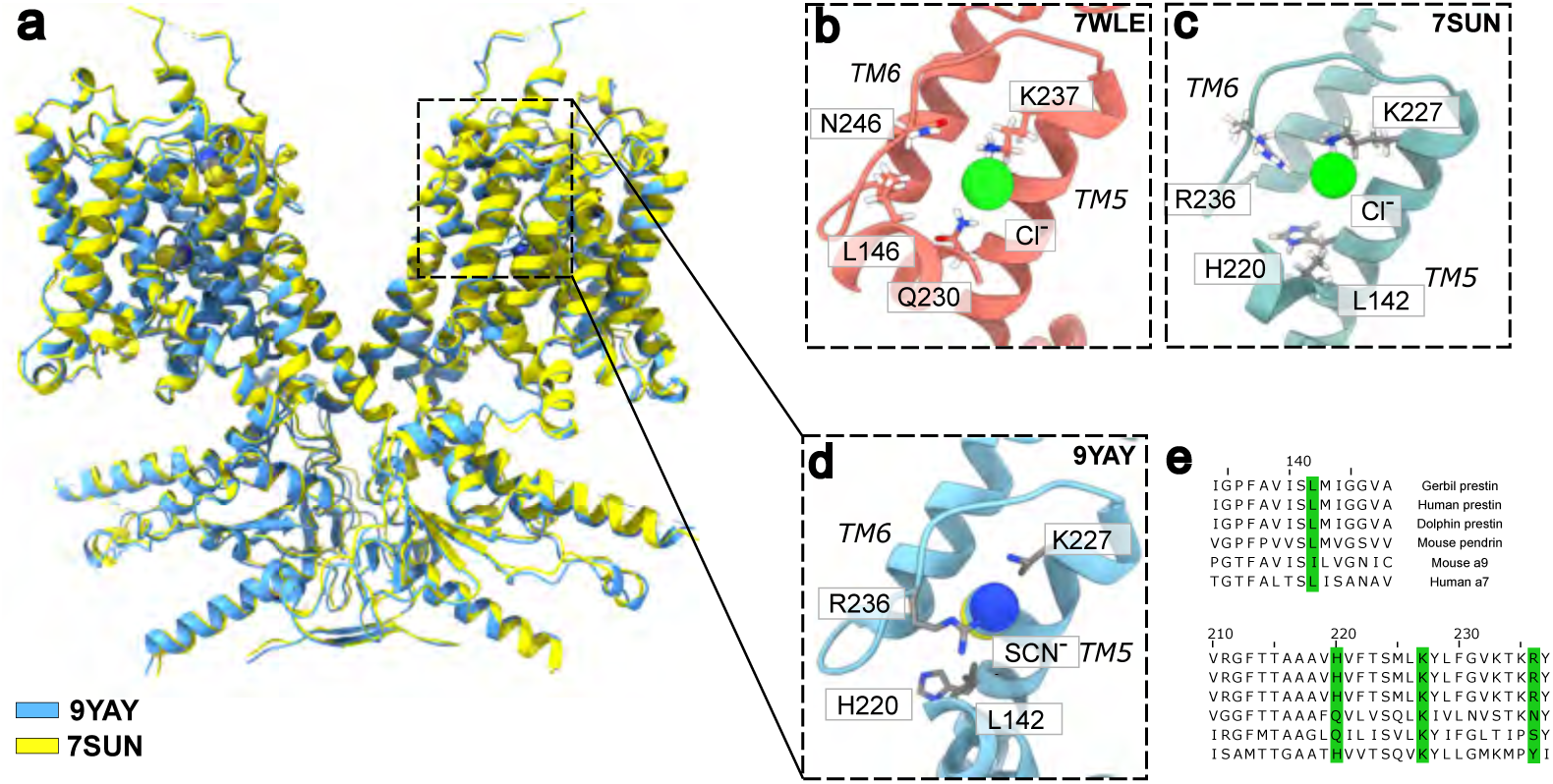
Anion binding in prestin and pendrin. Cryo-EM structure of prestin (PDB:9YAY) shows anion SCN^−^ binding in the IC and EC site. **a** New prestin cryo-EM structure (PDB:9YAY, 3.27 Å resolution, light blue), and gerbil prestin (PDB:7SUN, yellow). SCN^−^ ions are resolved at both the extracellular and intracellular binding sites of PDB:9YAY. **b** Pendrin extracellular anion binding site with Cl^−^ from MD simulation of OF pendrin. **c** Prestin extracellular anion binding site with Cl^−^ from MD simulation of prestin in the compact-2 state. **d** Prestin (PDB:9YAY) extracellular anion binding site with SCN^−^. **e** Sequence alignment of SLC26 family members with residues found in the EC site (gerbil prestin numbering).

### Cryo-EM structure confirms prestin EC anion site

We set out to confirm the presence of the EC anion site by solving the structure of gerbil prestin in the presence of the thiocyanate (SCN^−^) anion, where large currents have been observed ^37,38^. The cryo-EM density map of prestin in the presence of sodium thiocyanate was determined at an overall resolution of 3.27 Å according to the gold-standard Fourier shell correlation (FSC) and retrieved from 71,880 particles (Supplementary Information Figure S11). This structure in the presence of SCN^-^ (PDB:9YAY) most closely resembled our previous structure of prestin in the presence of Cl^−^ (PDB:7SUN; Figure 6a). In turn, both prestin structures (PDB:9YAY and PDB:7SUN) most closely resemble the intermediate structure of SLC26A9 (PDB:6RTF) ^14^, with RMSD values of 1.042 and 0.986 Å, respectively; additional structural details are described in Supplementary Information.

In addition to the canonical internal anion-binding pocket (described in Supplementary Information), we observed an additional density consistent with SCN^-^ near the extracellular surface of prestin, suggesting the presence of a potential external anion-interaction site. The density is positioned close to Lys227 and Arg236, with estimated interaction distances of approximately 3.5 Å and 3.8 Å, respectively, and is further located above the side chain of Leu142 at 3.7 Å (Supplementary Information Figure S7d). The external SCN^-^ ion is separated from the internally bound SCN^-^ in the canonical binding pocket by approximately 13.2 Å, indicating that the two ions occupy distinct positions along the transporter-like architecture of prestin.

In pendrin, residues Lys237 (corresponding to prestin Lys227, see sequence alignment in Figure 6e) and Gln230 (equivalent to prestin His220) have been identified as the external binding site ^20^ and therefore, we surmise, the equivalent binding pocket in prestin serves a similar purpose, collectively defining the extracellular anion binding site in prestin (Figure 6c, d). While it was not identified in any of the initial structures of prestin, we note a corresponding density in the 2.3-Å resolution electron density map of human prestin (PDB:7LGU) obtained by Ge et al. ^9^ that we believe represents a Cl^-^ anion (Supplementary Information Figure S7a). This density is located in the same location as the EC Cl^−^ density found in our MD simulations (Supplementary Information Figure S7c) and the EC SCN^-^ site in our cryo-EM structure (Supplementary Information Figure S7d).

A related feature has also been reported in SLC26A7, where iodide occupies the non-canonical site-2 (I2) pocket (external binding site) ^23^. In that structure, iodide interacts with Leu109, which is structurally equivalent to prestin Leu142. In SLC26A7 the iodide density lies below Leu109 at 3.8 Å, whereas in the present prestin structure the SCN^-^ density appears above Leu142 at 3.7 Å. The iodide external binding pocket in SLC26A7 is further defined by surrounding residues including Ala186, His189, Ala452, and Ala453, with His189 playing a particularly important role in stabilizing iodide and contributing to anion transport. Taken together, these comparisons indicate that the extracellular region of prestin centered on Lys227/Arg236 and Leu142 (Figure 6) correspond structurally to outer anion-interaction pockets described in other SLC26 transporters, suggesting that this region may represent a conserved second, extracellular anion-recognition site within the family.

### Conformational change connects anion binding sites in pendrin but not prestin

The prestin EC site matches the extracellular anion binding site described for the pendrin OF structure (PDB:7WLE) ^20^. In our simulation of OF pendrin, Cl^−^ binds intermittently to Lys237 and N246 (equivalent to Arg236 in prestin) and also Gln230 (Figure 6b and Supplementary Information Figure S7b), thus also broadly confirming the location of the EC site seen in the cryo-EM structure at the position of a double peak in the electron density ^20^. Closer inspection of the Cl^−^ density (Figure 6c and Supplementary Information Figure S9b) from the simulation of OF pendrin indicates that it extends from the EC site to the IC site and analysis of the ion trajectories confirmed that Cl^−^ ions entered the IC site from the extracellular compartment through the EC site. The visibility of continuous water density confirms that the IC site is accessible from the extracellular compartment but not from the intracellular one when pendrin is in the OF conformation (Figure 7a). In the IF conformation, the IC site in pendrin is accessible from the intracellular compartment and there is no connection between EC and IC sites visible (Figure 7b); the EC site may still be water accessible (Figure 7b) but possibly occluded to Cl^−^ (Supplementary Information Figure S9d). Taken together, the simulation data suggest that the IC site in pendrin appears to be the anion transport site that switches accessibility as part of the alternating access transport cycle and that only in the OF conformation, anions pass between the extracellular compartment and the IC site through the EC site.

**Figure 7.**
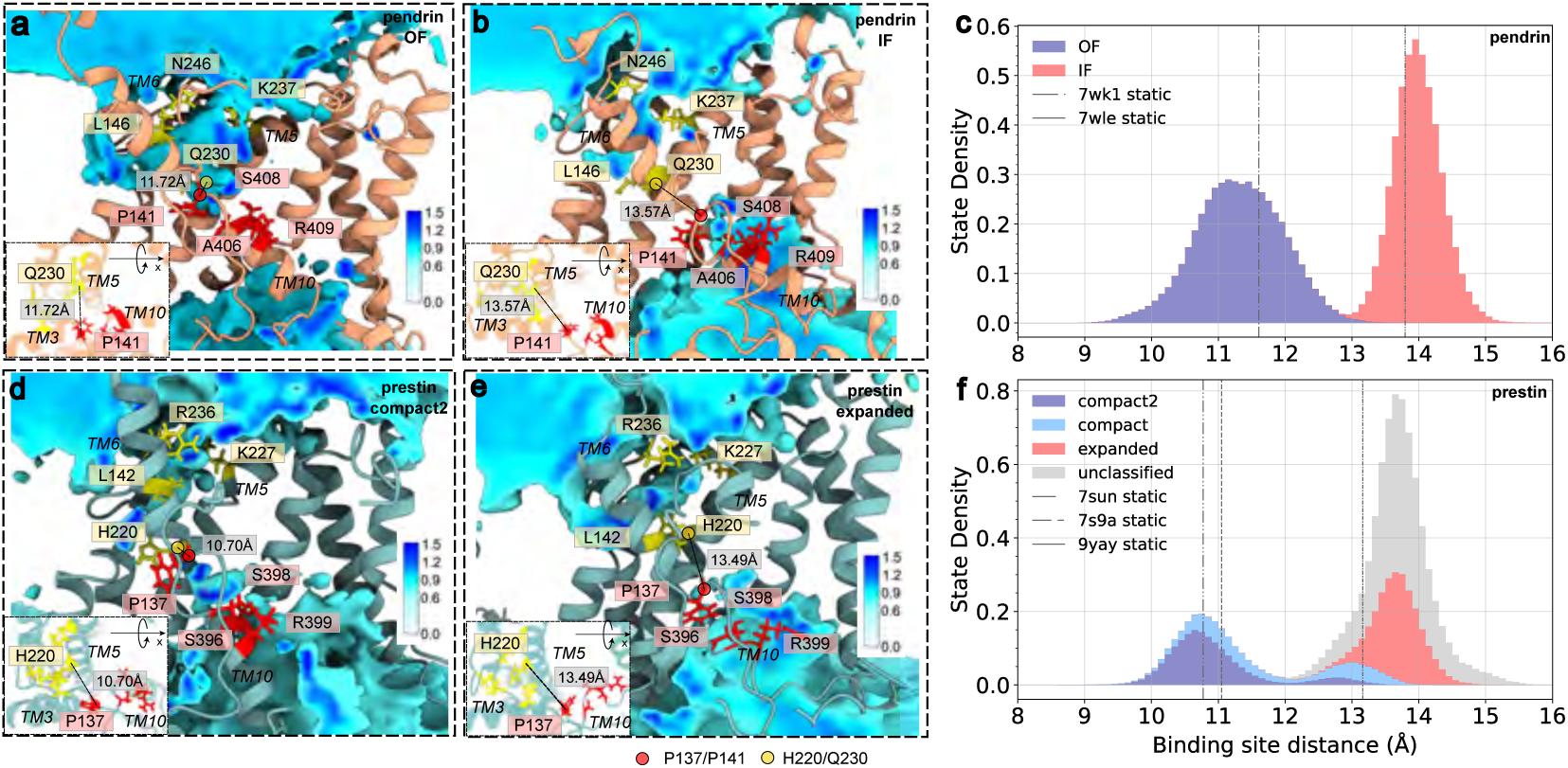
Conformation-dependent connectivity between anion binding sites in pendrin and prestin. **a** Water density in pendrin in the OF conformation, from MD simulation pendrin-outward. The water densities are scaled relative to the bulk water molarity (55.5 M) so that a value of 1 represents water density at standard conditions. Some helices were removed for clarity. Residues belonging to the canonical intracellular binding site in pendrin (Pro141, Ala406, Ser408, Arg409) or prestin (Pro137, Ser396, Ser398, Arg399) are labeled in red; residues in the extracellular binding site (pendrin: L146, Gln230, Asn246, Lys237) or (prestin: L142, His220, Arg236, Lys227) are labeled in yellow. The binding site distance *d*_EC,IC_ is indicated by a black dashed line between the highlighted C*_α_* atoms. *Insets* show a view of the binding site distance perpendicular to the connecting line. **b** Water density in pendrin in the IF conformation, from MD simulation pendrin-inward. **c** Probability distribution of the binding site distance *d*_EC,IC_ in pendrin from OF (blue) and IF (red) MD simulations. The site distance for the static experimental structures are indicated by the vertical lines. **d** Water density in prestin in the compact-2 conformation. **e** Water density in prestin in the expanded conformation. f Stacked probability distribution of *d*_EC,IC_ in prestin from all prestin MD simulations. Contributions to the total from simulation frames classified as compact-2 (blue), compact (light blue), expanded (red), and unclassified (gray) are indicated. Molecular images were produced with ChimeraX ^39^.

For prestin, however, no solvent-accessible pathway was observed between the EC site and the IC site, regardless of the conformation (Figure 7e, d and Supplementary Information Figure S8a). The IC site was water- and ion-accessible in the expanded (Figure 7e) and compact state (Supplementary Information Figure S8a) but somewhat less so in the compact-2 state as indicated by the broken water density in the simulations (Figure 7d). Unlike pendrin, the EC site appeared to be separated from the IC site, regardless of prestin’s conformation.

Because a solvent connection between EC and IC site was only observed when both sites were spatially adjacent, we computed the binding-site distance *d*_EC,IC_ between the C*_α_* atoms of the closest site residues (His220–Pro137 in prestin; Gln230–Pro141 in pendrin; see Figure 7). This distance clearly separates the OF and IF MD simulations of pendrin (Figure 7c). When *all* gerbil prestin simulations are analyzed in aggregate, a clear bi-modal distribution emerges (Figure 7f), which is consistent with our independent classification of the prestin simulations into contracted C (compact-2 and compact) and expanded E states. Furthermore, almost all our unclassified conformations show expanded-like *d*_EC,IC_ distances, indicating that the unclassified conformations are also mostly conformations resembling the expanded state, with only a small fraction of contracted-like conformations. The site distance correlates with the conformational state of the protomer (as seen in the projection of *d*_EC,IC_ on the *ζ*-*φ* order parameters, Supplementary Information Figure S10), with expanded-like conformations having larger distances (mode of the right peak of the distribution *d*_EC,IC_ = 13.8 Å) than contracted-like conformations (*d*_EC,IC_ = 10.9 Å).

Comparison of the MD simulations of pendrin and prestin indicates that only in pendrin the connection between the EC and IC sites depends on the conformational state. In pendrin, the canonical IC site is only accessible from the extracellular compartment in the OF state; in the IF state, only the IC site is accessible from the intracellular compartment. In prestin, the connection between the two anion sites is always interrupted, regardless of the conformational state and the IC anion binding is not accessible from the extracellular compartment for any of the observed conformations.

## Discussion

MD simulations of prestin initialized in the compact conformation, one of the two contracted conformations, typically transition on the microsecond timescale to an expanded conformation with a 150-Å^2^ area increase near the inner membrane leaflet. In the expanded state, prestin resembles the IF structure of pendrin, with the anion binding site accessible from the intracellular compartment. Prestin simulations also sample the stable contracted compact-2 conformation, which has the same cross-sectional area as the compact state but bears more structural similarity to OF pendrin. Our simulations and structural analyses support the view that the expanded state that we obtained represents a plausible and physiologically relevant conformation of prestin. Unlike previously reported expanded-like models, our expanded state conformation is not stabilized by exogenous ligands and instead emerges spontaneously from simulations initiated in the compact state. Its overall architecture closely resembles the IF conformation of pendrin. Existing experimental prestin structures ^9,10^ and MD-derived models ^34^ occupy positions along the same continuum between contracted and expanded states, as revealed by our order parameter analysis, further supporting the hypothesis ^26^ that prestin’s expansion/contraction cycle is analogous to part of the SLC26A4’s transporter cycle (Figure 8). Prestin may undergo conformational changes regardless of binding of Cl^−^ ions, based on the experimental findings that Cl^−^ is not strictly necessary for NLC but rather acts as an allosteric modulator, showing state-dependent Cl^−^ binding affinity ^12,13,40^, and the observation of contracted conformations in our simulations (compact and compact-2) without bound anions. Such transitions without an anion bound are likely forbidden for the transporter pendrin as these would lead to leak cycles that deplete the Cl^−^ ion gradient ^16^.

**Figure 8.**
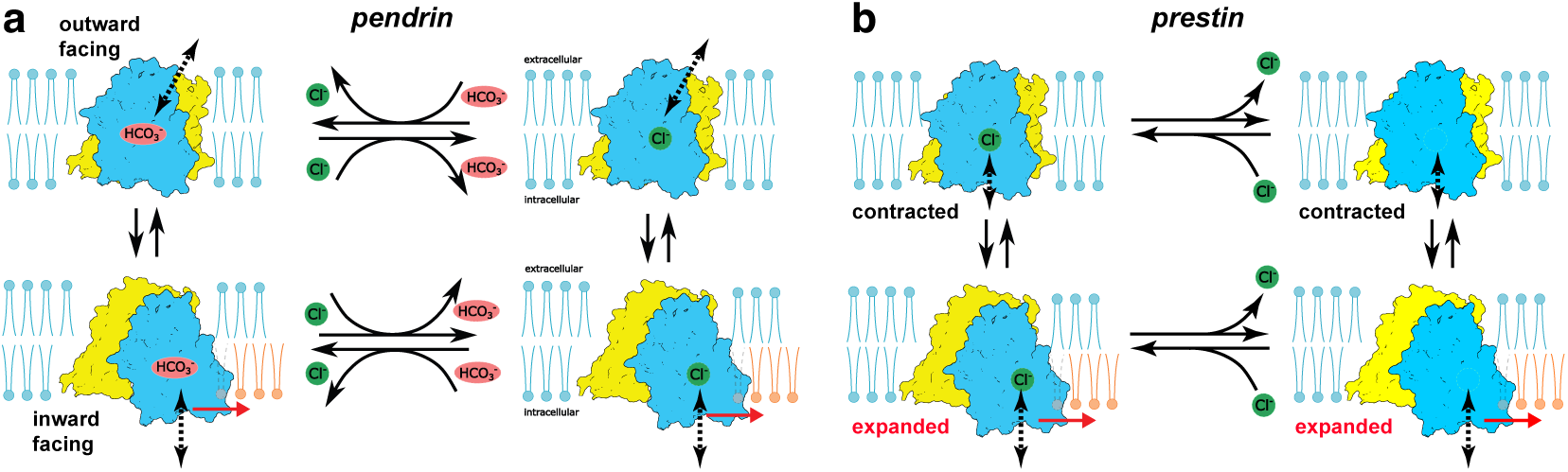
Comparison of conformational states of pendrin and prestin. **a** The alternating access transport cycle of pendrin includes outward facing (OF) and inward facing (IF) states with either Cl^−^ or HCO^−^ bound. The intracellular anion binding site is accessible from the extracellular compartment in the OF state (via the extracellular anion binding site, not shown) and from the intracellular compartment in the IF state (black dashed arrows). The core domain (blue) moves relative to the gate domain (yellow). In the IF conformation, pendrin’s cross-sectional area expands at the inner leaflet (red arrow) and displaces lipids. **b** Prestin exists in an equilibrium between contracted and expanded macrostates. The contracted macrostate comprises the compact state and the OF-like compact-2 state observed in our simulations. As in pendrin, the IF state expands at the inner leaflet due to rotation and vertical displacement of the core domain relative to the gate domain. In the contracted state, the anion binding site is not accessible from the extracellular compartment but may be accessible from the intracellular compartment (black dashed arrows). Prestin may undergo conformational changes regardless of binding of Cl^−^.

Our simulations show reproducible and rapid transitions from the initial compact conformation to an expanded conformation on the sub-microsecond timescale, consistently across gerbil and human prestin and across two independent MD engines (Anton 2 and GROMACS). Several independent lines of evidence indicate that this expanded conformation is physiologically relevant rather than a simulation artifact: it emerges spontaneously without inhibitory ligands, closely resembles the IF conformation of pendrin ^20^, lies along the same compact-to-expanded path as the sulfate- and salicylate-stabilized “expanded-like” prestin structures ^9,10^, and resembles the expanded state independently identified in equilibrium MD of human prestin by Kuwabara et al. ^34^ . The convergence of two simulation studies—using different starting structures, MD engines, and membrane and ionic conditions—together with multiple experimental structures on the same expanded endpoint suggests that this conformation is a physiologically relevant model for the expanded state of prestin.

A clear limitation is that our equilibrium simulations sample only the forward compact-to-expanded relaxation and never a reverse expanded-to-contracted transition. This lack of reversibility appears to be a common feature of unbiased equilibrium MD simulations of prestin: Kuwabara et al. ^34^ likewise populated both compact and expanded states in aggregate but did not observe reversible interconversion within trajectories. Reversibility of prestin’s expansion/contraction is in any case well established functionally by its bidirectional, voltage-dependent NLC and electromotility, and structurally by the capture of prestin in both contracted and expanded-like states. The absence of the reverse transition in our trajectories therefore reflects the simulation timescale, rather than a true thermodynamic irreversibility. Several non-exclusive factors may additionally bias the simulations toward the expanded state: current MD force fields may not perfectly reproduce the relative energetics of the contracted and expanded macrostates; the cryo-EM compact structure may represent a state stabilized by detergent or nanodisc conditions rather than the resting state at 0 mV; and our simplified POPC/cholesterol membrane lacks the specialized lipids of the native OHC membrane that could further stabilize the contracted state *in vivo*. Consistent with the compact basin being metastable rather than absent, a subset of trajectories remained compact for their full duration (Supplementary Information Table S1) and two trajectories instead sampled the longer-lived compact-2 state resembling OF pendrin. So far, the majority of the structurally studied SLC26 family members have been observed to adopt an IF conformation ^8–10,14,19–24^ (and several other members of the extended family including UraA and Slc26Dg ^41–43^). Only pendrin (SLC26A4) was also observed in the OF conformation, in the presence of 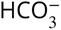 and Cl^−^, natural substrates for its exchange mechanism^20^. In our simulations we also observe a majority of conformations of prestin with the canonical anion binding site being accessible from the intracellular side (expanded, compact) and fewer ones resembling OF transporter conformations (compact-2), which qualitatively matches the sampling of conformations across cryo-EM experiments of the many different SLC26 family members. Thus, the simulation conditions we used are likely more similar than dissimilar to those in the cryo-EM experiments as the proportion of structures in the IF and OF forms shows the same tendency to favor IF conformations.

Prestin’s (and pendrin’s) expansion results from a non-uniform and preferential expansion of the TM domains closer to the inner leaflet. The estimated expansion of the cross-sectional area of a single protomer is about 150 Å^2^ or 8.8%, comparable to previous estimates ^9,10,34^ and directly compatible with estimates for the expansion of prestin that is required to support the experimentally observed voltage-induced elongation of OHCs ^6,44^. As we showed here and in previous work ^45^, neutralization of specific charged residues closer to the inner leaflet results in a reduction in *z* (the estimate of the charge carried by a single prestin motor), which is associated with charge movement relative to the membrane potential. The localization of these neutralizing mutations to the region where domain movements have the largest effect in MD substantiates our simulation data. While a full mechanistic interpretation of the observed inner leaflet expansion is difficult, these data have parallels in the membrane bending model initially proposed by Raphael and Brownell where expansion occurs at the inner leaflet and is brought about by depolarization induced alignment of the membrane dipoles ^7^. Irrespective, the implications of asymmetric membrane distortion to produce force are significant and have been modeled and observed in other mechanosensitive channels including piezo1 and MscL ^46,47^; coarse-grained MD simulations of prestin suggested that such asymmetries may facilitate long-range coupling between dimers in the membrane ^48^. In addition to lipid composition, asymmetric membrane distortion would also suggest roles for membrane lipid transporters in tuning mechanical responses. Expansion at the inner leaflet appears to be the first step by which prestin couples changes in the membrane potential to the membrane and further work is needed to understand how these changes in the lipid bilayer lead to OHC elongation or force generation.

Despite uncertainties in the comparative ability of prestin and pendrin to transport anions or function as a motor protein, prestin closely resembles pendrin in many aspects. The extracellular chloride-binding site appears structurally conserved in prestin, pendrin, and likely other SLC26 family members. We previously suggested that prestin might have more than one anion binding site ^13,49,50^. While its exact role has yet to be determined, we note that neutralization of Lys227 to Gln in the prestin EC site results in a reduction in macroscopic whole-cell currents in SCN^−^, along with effects on NLC with a hyperpolarizing shift of *V*_ℎ_ in Cl^− 38,45^, thus indicating the EC site may exert an effect on prestin’s function.

We found that in MD simulations of pendrin in the OF state, the canonical intracellular anion binding site was accessible from the extracellular compartment, with a solvated pathway leading through the extracellular site. In the IF conformation, the intracellular anion binding site is accessible from the intracellular compartment and it is not connected to the EC site. In the context of the alternating access mechanism, this finding implies that at least in pendrin, the intracellular anion site is the transport site because of its conformation-dependent accessibility. The extracellular site may play a role in electrostatically attracting anions in the OF conformation and thus kinetically enhance transport ^16,17^. In prestin, no similar pathway connecting EC and IC anion sites was found, neither in the expanded or compact conformations nor in the compact-2 conformation, even though the latter structurally most closely resembles OF pendrin. We hypothesize that this compact-2 conformation may represent a state near the end point of the physiological voltage-driven eM work cycle of prestin, during which no Cl^−^ ions are transported across the membrane, consistent with the observed lack of a pathway to the EC site. Mammalian prestin has been shown to also function as a slow 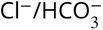 antiporter ^51^ and possibly as an oxalate/Cl^−^ antiporter ^45^ and therefore the compact-2 conformation may also represent an intermediate along a rare full transporter-like conformational cycle of prestin.

Our simulations reveal that the two prestin protomers within the homodimer behave independently on the microsecond timescale. In several trajectories, one protomer transitions to the OF (compact-2) conformation while the other remains in the IF (expanded) conformation within the same dimer. While we did not observe such behavior with pendrin in simulations, its cryo-EM structures showed conformational heterogeneity including concurrent OF and IF protomers ^20^. While these data seemingly contradict the findings of Detro-Dassen et al. ^52^ that suggested cooperativity, the findings do not discount one protomer influencing the other while still moving in opposing directions.

The conformational changes we observe in our simulations of prestin map onto a partial transition of pendrin, and although we do not simulate pendrin’s full transport cycle, comparison of endpoint structures indicates that the principal moving element, namely the elevator-like transition of the core domain relative to the gate domain, is shared between the two proteins (Figure 8). Pendrin also undergoes a comparable magnitude of cross-sectional area expansion when transitioning from its OF to IF state, mirroring the expansion we observe in prestin between contracted and expanded macrostates.

Moreover, electrophysiological measurements presented here and by Kuwabara et al. ^36^ demonstrate that pendrin presents nonlinear capacitance (NLC), indicating that, like prestin, its area expansion is necessarily coupled to changes in membrane potential, and it could conceivably also act as an electromechanical transducer, especially if its *V*_ℎ_ could be right-shifted.

Despite similarities, key differences distinguish prestin from pendrin. In our simulations, pendrin starting in either OF or IF conformations remained in these states with no transitions between them. In contrast, prestin showed rapid movements on a sub-µs timescale from the intermediate (compact) to IF (expanded) or more OF-like (compact-2) conformations. While these rapid transitions in prestin may reflect the energy landscape of the intermediate structure in MD simulations, these data hint that prestin’s role as a fast electromechanical coupler may stem in part from its intrinsic rapid structural fluctuations, which are not shared with pendrin. In this context it is important to note that OHC eM has been observed to occur at ultrasonic frequencies ^27–33^, but it still remains unclear what in prestin’s intrinsic structure poises it to move at such speeds.

## Methods

### Multiple sequence alignments

A multiple sequence alignment of prestin and pendrin homologs was generated using Jalview web services ^53^. Sequences were obtained from UniProt, including gerbil prestin (Q9JKQ2), human prestin (P58743), dolphin prestin (D7PC76), mouse pendrin (Q9R155), and mouse Slc26a7 (Q8R2Z3).

### Prestin expression and purification

Full-length gerbil prestin (*Meriones unguiculatus*, GenBank accession number AF230376) was purified from a tetracycline-inducible stable HEK293 cell line (clone 16C) as previously described ^8^. Briefly, the gerbil prestin cDNA, fused at its C terminus to enhanced yellow fluorescent protein (EYFP), was subcloned into the pcDNA4/TO/myc-HisC vector, enabling purification by Ni-NTA affinity chromatography. The construct was kindly provided by J. Zheng and P. Dallos. Cells were cultured in DMEM supplemented with 1 mM L-glutamine, 100 U ml^-1^ penicillin/streptomycin, 10% fetal bovine serum (FBS), and 1 mM sodium pyruvate. Prestin expression was maintained under selective pressure with 4 µg ml^-1^ blasticidin and 130 µg ml^-1^ zeocin. Protein expression was induced by addition of 1 µg ml^-1^ tetracycline for 48 h. Cell pellets harvested from 20 T175 flasks were collected by centrifugation at 1,000 g for 10 min, washed in PBS, and resuspended at 5 ml buffer per gram of pellet in resuspension buffer (25 mM HEPES pH 7.4, 200 mM NaCl, 5% glycerol, 2 mM CaCl_2_, 10 *μ*g⋅ml^-1^ DNase I, and protease inhibitor cocktail [Roche, EDTA-free]). Solubilization was performed by direct addition of digitonin powder (final concentration 2% w/v), followed by incubation with gentle agitation for 1.5 h at 4 °C. The lysate was clarified by ultracentrifugation (160,000 g, 50 min, Beckman L90-XP, 50.2 Ti rotor), and the supernatant was filtered through a 0.45 µm membrane. The supernatant was supplemented with 10 mM imidazole and incubated with 1 ml of Ni-NTA resin (Qiagen), pre-equilibrated in buffer containing 0.02% glyco-diosgenin (GDN), for 2 h at 4 °C with gentle agitation. The resin was collected and sequentially washed with high-salt Buffer A (25 mM HEPES pH 7.4, 500 mM NaCl, 5% glycerol, 0.02% GDN) and Buffer B (25 mM HEPES pH 7.4, 200 mM NaCl, 30 mM imidazole, 5% glycerol, 0.02% GDN). Bound protein was eluted in Buffer C (25 mM HEPES pH 7.4, 200 mM NaCl, 250 mM imidazole, 5% glycerol, 0.02% GDN). The eluate (∼3 ml) was concentrated to 600 µl, filtered through a 0.22 µm membrane, and subjected to size-exclusion chromatography (Superdex 200 Increase 10/300 GL, Shimadzu FPLC) pre-equilibrated in gel-filtration buffer (10 mM HEPES pH 7.4, 200 mM NaCl, 0.02% GDN). Fluorescent and A280 peaks corresponding to prestin were pooled, concentrated using a 100 kDa cutoff Amicon Ultra centrifugal filter, and used for cryo-EM grid freezing.

### Sample preparation and cryo-EM data acquisition

Aliquots of purified prestin (∼3 mg⋅ml^-1^) were supplemented with 40 mM sodium thiocyanate (SCN^-^) and incubated for 10 min prior to grid freezing. Three microliters of protein solution were applied to glow-discharged Quantifoil R 1.2/1.3 gold grids (300 mesh). Grids were blotted for 0–5 s under 100% humidity at 8 °C and plunge-frozen into liquid ethane using a Vitrobot Mark IV (FEI).

Cryo-EM data were collected on a Titan Krios G2 transmission electron microscope (Thermo Fisher Scientific) operating at 300 kV and equipped with a K3 Summit direct electron detector (Gatan) and a 20-eV slit-width energy filter. A total of 5076 super-resolution movies were recorded with SerialEM (v3.8), using a nominal magnification of 81,000× (corresponding to 0.534 Å physical pixel size). Movies comprised 40 frames, acquired at a dose rate of 19.3 e^-^⋅pix^-1^⋅s^-1^, with a total exposure time of 3.2 s and cumulative dose of ≈54 e^-^⋅Å^-2^. The nominal defocus range was set between –2.0 and –2.5 µm.

### Data processing

All image processing was performed in CryoSPARC (v4.3.1) ^54^. Movie frames were gain-normalized, aligned, and dose-weighted using patch motion correction with a 5 × 5 patch grid and a binning factor of 2. CTF parameters were estimated using patch CTF estimation (multi-micrograph mode) on non-dose-weighted sums. Template-based particle picking was performed using 2D projections generated from the previously determined prestin structure (PDB:7SUN) low-pass filtered to 20 Å. This approach yielded approximately ∼250,000 particles. Multiple rounds of 2D classification were carried out to remove false positives and damaged particles, resulting in a curated dataset. Subsequent ab-initio 3D reconstruction and heterogeneous refinement were used to eliminate remaining heterogeneous classes. A final subset of 71,880 particles was subjected to non-uniform refinement with C2 symmetry imposed, yielding a reconstruction at 3.27 Å resolution (gold-standard Fourier shell correlation at 0.143).

### Model building and refinement

The cryo-EM structure of gerbil SLC26A5 (PDB:7SUN) was initially rigid-body docked into the prestin density map using UCSF Chimera (v1.12) ^39^. Manual rebuilding was performed in COOT (v0.9.8.1) ^55^ with sequence assignment guided by bulky side chains (Phe, Tyr, Trp, Arg) and secondary structure predictions. Iterative cycles of real-space refinement in Phenix (^56,57^ v1.21.2), combined with manual adjustments in COOT ^55^, were used to optimize model geometry. Refinement was carried out with secondary structure restraints and stereochemical constraints. Model validation was performed using MolProbity, and refinement statistics are summarized in Supplementary Information Table S6. Unmodeled rod-like densities were present on the final map near the canonical chloride-binding site and an external binding site. Thiocyanate ions were placed into these densities. The structure was deposited in the PDB submission D_1000300147 with the following accession codes: PDB:9YAY, extended PDB ID pdb_00009YAY, and EMD-72742.

### Electrophysiology

Prestin mutants were generated using QuickChange II site-directed mutagenesis kit (Stratagene) with gerbil prestin-YFP in pEYFPN1 vector (Clontech) as a template as previously described ^38^. All mutations were confirmed by DNA sequencing. Transient transfection of constructs into CHO cells was done using lipofectamine 2000 (ThermoFisher Scientific) according to the manufacturer’s instructions, in 24-well plates ^38^. Cells were recorded 48–72 h after transfection in whole-cell configuration at room temperature using an Axon 200B amplifier (Molecular Devices) ^38^. Pipette resistance was 3–5 MΩ. For pendrin transfections we subcloned human pendrin cDNA purchased from Addgene (pDONR221_SLC26A4 (Plasmid #132032) Addgene.Org) into pEYFPN1 as with prestin. Transient transfection conditions and recordings were as described for prestin. Bath solution: TEA 20 mM, CsCl 20 mM, CoCl_2_ 2 mM, MgCl_2_ 1.47 mM, HEPES 10 mM, NaCl 99.2 mM, and CaCl_2_ 2 mM, pH 7.2. Pipette solution: CsCl 140 mM, EGTA 10 mM, MgCl_2_ 2 mM, and HEPES 10 mM, pH 7.2. Osmolarity was adjusted to 300±2 mOsm with dextrose ^38^. For recording in sulfate-based solution, the bath solution contained: CsCl 10 mM, Na_2_SO_4_ 95 mM, MgSO_4_ 5 mM, and Tris-OH 20 mM, pH adjusted to 7.2 using methanesulfonic acid, and the pipette solution contained: CsCl 10 mM, EGTA 10 mM, Cs_2_SO_4_ 95 mM, MgCl_2_ 2 mM, and HEPES 10 mM, pH 7.2^33,38^. Command delivery and data collections were carried out with a Windows-based whole-cell voltage clamp program, jClamp (Scisoft, East Haven, CT, www.scisoftco.com), using a NI USB DAQ device (National Instruments). A continuous high-resolution 2-sine voltage command was used ^58,59^. Capacitance data were fitted to the first derivative of a two-state Boltzmann function

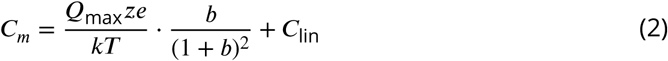

where

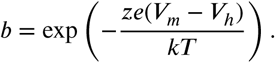

*Q*_max_ is the maximum nonlinear charge transfer, *V*_ℎ_ the voltage at peak capacitance or half-maximal nonlinear charge transfer, *V_m_* the membrane potential, *C*_lin_ linear capacitance, *z* the unitary charge, *e* the electron charge, *k* the Boltzmann constant, and *T* the absolute temperature. *Q*_sp_ the specific charge density, is the total charge moved (*Q*_max_) normalized to *C*_lin_. The effects of different parameters of NLC were evaluated by Student’s *t*-test. Data were presented as the mean ± SEM.

### All-atom molecular dynamics simulations

We performed MD simulations of prestin (from gerbil and human) and pendrin (from mouse, in IF and OF conformations) under equivalent conditions. The simulations are summarized in Table 1 and details are provided in the following sections.

### Membrane protein system preparation

In order to perform all-atom, explicit solvent MD simulations we constructed a model of gerbil prestin from the cryo-EM structure (PDB:7SUN ^8^) and the AlphaFold2^60^ gerbil prestin model (Q9JKQ2 S26A5MERUN) as described in Bai et al. ^33^ using the following approach: The unresolved gap in the structure (PDB:7SUN) at residues 159-164 was filled with the corresponding residues in the AlphaFold structure after the two models had been superimposed at residues 138-158 and 165-194. Residues 581-613 in the STAS domain, which are not resolved in any structures, predicted to be unstructured, and not reliably modeled by AlphaFold, were omitted from our simulation system. The resulting model of wildtype (WT) gerbil prestin was set up in a 1-palmitoyl-2-oleoyl-sn-glycero- 3-phosphocholine (POPC) and cholesterol bilayer with a 9:1 ratio and surrounding solvent with a free NaCl concentration of 150 mM using the CHARMM-GUI V1.7 web interface ^61,62^.

Simulations of human prestin (*Homo sapiens*) were based on the cryo-EM structure (PDB:7LGU) ^9^, which did not require any loop-filling, except that its unstructured domain was also omitted. The bound Cl^−^ ion was removed to match the situation in the gerbil prestin structure. In all other aspects the same set-up and simulation protocol as for gerbil prestin was followed.

We set up simulation systems for mouse pendrin (*Mus musculus*) using the cryo-EM structures for the IF (PDB:7WK1) and OF (PDB:7WLE) conformations ^20^ in the same manner as for prestin.

All simulations used the CHARMM36 force field ^63^ for proteins ^64,65^, ions, and lipids ^66^ as well as the CHARMM TIP3P model for water. Titratable residues were simulated in their default protonation states at pH 7 as predicted by PROPKA 3.1^67^. System sizes ranged from 208096 to 247744 (see Table 1 for a list of all simulations together with the IDs by which they are referred to in the text).

Systems were energy minimized, relaxed, and equilibrated using GROMACS ^68,69^ at temperature 296 K and pressure 1 bar by following the standard CHARMM-GUI protocol with an initial energy minimization stage and six stages of equilibration during which harmonic restraints on protein and lipids were successively reduced to zero ^61^.

### MD simulations on Anton 2

Production runs of prestin were performed on the Anton 2 supercomputer, a special-purpose computer for molecular MD simulations of biomolecules ^70^. The CHARMM36 force field was used for proteins and ions, and the TIP3P model was used for water, as provided on Anton 2. The RESPA algorithm ^71^ was used to integrate the long-range nonbonded forces every 7.5 fs and short-range nonbonded and bonded forces every 2.5 fs. Long-range electrostatic forces were calculated using the *k*-Gaussian split Ewald method ^72^. The lengths of bonds to hydrogen atoms were held fixed using the nSHAKE algorithm. The simulations were run at 296 K and 1 atm using the Nosé-Hoover chain thermostat ^73^ and the Martyna-Tobias-Klein barostat ^74^, respectively. The RESPA algorithm and temperature and pressure controls were implemented using the multigrator scheme with a time step of 2.5 fs ^75^. Temperatures were maintained using Nosé-Hoover chains with time constants *τ_T_* = 0.042 s. Semi-isotropic pressure coupling using Nosé-Hoover chains kept the system at 1 atm across both temperature groups with time constants *τ_P_* = 0.042 s. Coordinates were written to trajectory files every 240 ps.

We were interested in resolving the initial steps of the prestin-Anton-0mV simulation, and therefore we restarted the exact same trajectory from the initial conditions with a write-out interval of 1.2 ps and total run time of 10 ns. The specific architecture of

Anton 2^70^ guarantees that these two trajectories are binary identical. This trajectory is not listed with the other production simulations in Table 1.

### GROMACS MD simulations

Production runs of prestin were performed on Arizona State University’s sol supercomputer ^76^ and local workstations with GROMACS version 2024.2^68,69^. Run parameters were chosen to closely match the Anton 2 simulations with adjustments to account for different MD codes and hardware. The stochastic velocity rescaling thermostat ^77^ was used with a time constant of 1 ps and three separate temperature-coupling groups for protein, lipids, and solvent. The Parrinello–Rahman barostat with a time constant of 5 ps and a compressibility 4.5 × 10^-5^ bar^-1^ was used for semi-isotropic pressure coupling ^78^. Coulomb interactions were calculated with the fast-smooth particle-mesh Ewald (SPME) method ^79^ with an initial real-space cutoff of 1.2 nm, which was optimized by the GRO- MACS GPU code at run time, and interactions beyond the cutoff were calculated in reciprocal space with a fast Fourier transform on a grid with 0.12 nm spacing and fourth-order spline interpolation. The Lennard–Jones forces were switched smoothly to zero between 1.0 and 1.2 nm and the potential was shifted over the whole range and decreased to zero at the cutoff. Bonds to hydrogen atoms were converted to rigid holonomic constraints with the P-LINCS algorithm ^80^, except for water, in which all bonds were constrained with the analytical SETTLE algorithm ^81^. The classical equations of motion were integrated with the leapfrog algorithm with a time step of 2 fs. The initial production runs of pendrin were performed on the Yale supercomputing cluster with GROMACS 2021.6 and subsequent repeats of pendrin simulation were performed on the Arizona State University’s sol supercomputer with GROMACS 2024.2^68,69^. These simulations followed the same protocol used for prestin, except that a Nosé-Hoover thermostat ^82,83^ was employed for the initial pendrin simulations; both thermostat algorithms sample the correct statistical mechanical ensemble, so the specific choice does not affect simulation outcomes.

### Analysis of MD simulations

#### Structural similarity and clustering

Structural similarity was assessed with the root mean square distance (RMSD) using the fast QCP algorithm ^84^ in MDAnalysis ^85^. All simulations were stable as judged by the C*_α_* RMSD of the transmembrane helices of each protomer (Supplementary Information Figure S12). Although this C*_α_* RMSD hints at conformational changes, it is too coarse a measure to detect relative domain movements reliably. We therefore employed a different RMSD-based quantity for the majority of our structural analysis: Because we are primarily interested in domain movements in prestin and structurally similar SLC26 transporters, we calculated the C*_α_* RMSD of the core domain of each protomer (gerbil prestin residues 76–155, 167–196, and 339–426, mouse pendrin residues 80–159, 177–206, and 349–436) after optimal structural superposition on the gate domain (prestin residues 207–307 and 437–504, pendrin residues 217–317 and 447–514). This measure was found to clearly describe the large conformational domain movements that we observed in our simulations and that are apparent by comparing pendrin in its IF and OF conformation. We used this “core-RMSD” to compare all simulation frames to the pendrin IF reference conformation (PDB:7WK1).

To characterize the conformational states of prestin, we applied the GROMOS clustering algorithm ^86^ with the core-RMSD after superposition on the gate domain as the distance measure between structures. After clustering protomer B in the prestin-Anton0mV simulation with a cutoff of 2.5 Å, the centroid of the largest cluster, the frame at 3.372 µs, was taken to represent the expanded state. The conformation of the compact prestin structure (PDB:7SUN) after the initial energy minimization and equilibration steps was chosen to represent the experimentally determined compact state. Simulation prestingmx-1 sampled a new region of conformational space near the pendrin OF structure and GROMOS clustering of protomer B in prestin-gmx-1 with a 1.5 Å cutoff yielded a representative conformation for the compact-2 state as the centroid of the largest cluster; the smaller cutoff was chosen to better differentiate a homogeneous structural ensemble.

A structure-based state classifier was constructed to assign the conformation of a protomer in a trajectory frame to one of the three states (compact, compact-2, expanded). For each simulation frame, pairwise core-domain RMSDs were computed against the expanded, compact, and compact-2 reference structures. A protomer was then assigned to the state with the lowest RMSD, provided that the minimum value was ≤ 3.0 Å. Snapshots whose RMSDs exceeded 3.0 Å for all three references were designated unclassified (see Supplementary Information Figure S2).

### Bayesian rate estimate

The transition rate *k* was estimated following the method of Ensign and Pande ^35^ using Bayesian inference for a single exponential waiting time distribution with a uniform prior. In brief, for each trajectory *i* in an ensemble of *M* trajectories starting in the compact state, the time *t_i_* until it reaches the expanded state for the first time is recorded (Supplementary Information Table S1) as well as the total number *n* of observed transition events. If no transition occurs, then *t_i_* is set to the total trajectory length. The total time for observing the *n* transitions is 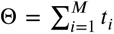. The maximum a-posteriori estimate for the rate is then *k* = *n*∕Θ and the posterior distribution is a Gamma distribution 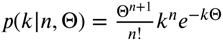.

### Structural order parameters

We describe the motions of the core domain relative to the stable gate domain in terms of rigid body motions relative to a reference structure, the IF pendrin structure (PDB:7WK1). Order parameters *ζ* and *φ* were computed for each protomer after optimal structural superposition of the protomer’s gate domain on the gate domain of the reference (as described above for the RMSD) following Okazaki et al. ^87^ with specific changes that increased robustness: The translational displacement vector between the centers of mass of the prestin and the reference (IF pendrin) core domains was calculated and the order parameter *ζ* was defined as its projection on the *z*-axis

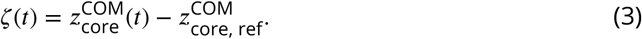

The translational component was then removed from both core domains. The rotation matrix R for optimally superimposing the core domain on the core domain of the reference structure was obtained using the MDAnalysis `rotation_matrix()` function ^85,88^ and its rotation angle

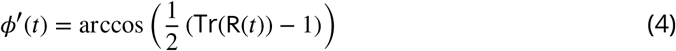

together with its rotation axis **n̂**(*t*) were extracted using `rotation_from_matrix()` ^85^. The axis is not returned in a fixed reference frame and because the pairs (*φ*′, **n̂**) and (−*φ*′, −**n̂**) describe the same rotation, the sign of the angle is undefined; Okazaki et al. ^87^ used *φ*′ as the order parameter but this approach introduces ambiguity, especially for angles near 0. As described in detail in the Supplementary Information, we sought to obtain a robust *signed φ* by determining the sign of each *φ*′ by comparison with the rotation about a fixed reference axis **m̂**, chosen as the axis that rotates the core domain of OF pendrin (PDB:7WLE) onto the core domain of IF pendrin (PDB:7WK1) and oriented such that this rotation angle is negative; consequently OF pendrin (PDB:7WLE) and compact prestin (PDB:7SUN) are recorded as negative. The values of *ζ* = 0 and *φ* = 0 correspond to the IF pendrin reference structure (PDB:7WK1).

The rotation of each trajectory frame was described by the unit quaternion 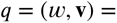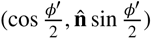 with *w* ≥ 0 and projected onto a rotation about **m̂**, i.e., we determined the signed angle *θ* for which the rotation by *θ* about **m̂** is closest to the rotation described by

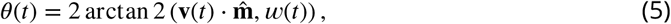

with the two-argument arctan 2 function (see Supplementary Information equation (S5) for the derivation). The projection *θ* is signed but noisy, whereas *φ*′ is robust and fully quantifies the magnitude of the conformational change, even when the projection axis **m̂** and the true rotation axis **n̂** are far from parallel. We therefore combined the two: with 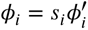 and sign states *s_i_* ∈ {−1, +1} for the *N* discrete time frames *i* of a trajectory, the sign path {*s_i_*} was chosen to minimize the objective function

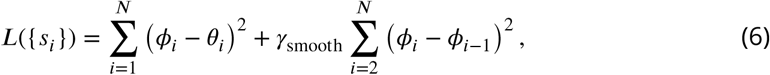

where the first term enforces agreement with the projected angle *θ_i_* = *θ*(*t_i_*) at time *t_i_* and the second term penalizes abrupt frame-to-frame changes that would imply unphysically fast conformational changes. Because each frame has only two sign states, the global minimum was obtained exactly by dynamic programming, and because only signs are assigned, the magnitude of the rotation angle is preserved, thus generating a robust *φ*(*t*) timeseries for large and small rotation angles (Supplementary Information Figure S13). For the smoothness hyperparameter, we used *γ*_smooth_ = 1000, the smallest value that suppressed all unrealistic sign jumps (Supplementary Information Figure S14).

### Area analysis

The protein area profiles (cross-sectional area of slices perpendicular to the z-axis) *A*(*z*) were calculated for all-atom systems using periodic 2D Voronoi tessellation. The key idea of this method is to use the boundary between protein and membrane or solvent to define the area that the protein covers in the native environment. In brief, for each trajectory frame at time *t* a single-frame protein area profile *A*(*t*, *z*) was calculated by the following steps: 1. The whole trajectory was superimposed onto the center of mass of the bilayer. 2. Atom positions were collected for slices along the z-axis at a spacing of 0.5 Å. 3. Voronoi tessellation was applied to the periodically replicated 2D atomic positions within each slice (using SciPy ^89^) and each atom was assigned to a Voronoi cell (see Supplementary Information Figure S15 for an example of a tessellated slice). 4. The areas of all Voronoi cells of protein atoms were calculated (using the shapely package from https://github.com/shapely/shapely) and summed to yield the area for the slice at position *z* and time *t*. Profiles were time-averaged to produce the average profile *A*(*z*), together with the corresponding standard error of the mean. Alternatively, we associate the maximum area at the peak of the profile with the trajectory frame *t*, *A*(*t*) = max*_z_ A*(*t*, *z*). The algorithm was implemented in the Python package Protein_Area whose source code is available under the open source GNU General Public Licenses v2 (or higher) at https://github.com/Becksteinlab/Protein_Area. Additional details and limitations of the algorithm and the package can be found in the Supplementary Information. Our approach enables us to directly calculate the full area profile of a membrane protein from an all-atom MD simulation.

In the simulations, each protomer in the dimer appears to change conformation independently (see Supplementary Information Figure S1) and we observe dimers with homogeneous states (C-C and E-E, where the contracted state C contains conformations classified as compact and compact-2, and E consists of expanded conformations) and heterogeneous states (C-E and E-C, but because the protomers are sequence-identical and thus functionally indistinguishable these two permutations are equivalent and so we consider them as just C-E). In order to associate the in-membrane expansion with a defined conformation we assume a simple linear model (equation (1)) in the number of protomers *n* in the expanded conformation (i.e., 0 for C/C, 1 for C/E, or 2 for E/E), whereby each of the two protomers contributes to the expansion independently and additively. We used SciPy ^89^ to perform a least-squares fit of all classified trajectory frames to the linear model in order to estimate Δ*A* and *A*_0_.

The area contribution *A_i_*(*t*) of one protomer *i* (protein chain A or B) in state *s_i_* (E or C) to the total measured dimer area *A*_dimer_ (*t*) = *A*_A_(*t*)+*A*_B_(*t*) at time *t* is calculated proportionally from the fitted linear model and the dimer area: if *s*_A_ = *s*_B_ (i.e., C-C or E-E) then 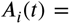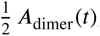; for a mixed conformational state (C-E), 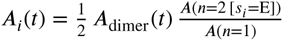, where the Iverson bracket [*s_i_* = E] equals 1 if protomer *i* is classified as being in the exanded state and 0 otherwise. Only trajectory frames with both protomers being classified as one of E or C were analyzed.

### Binding analysis

We analyze binding to the extracellular (EC) anion binding site in prestin with an approach similar to our previous analysis of the canonical intracellular (IC) site ^33^ although we first had to identify the residues forming the site from the MD simulations. We started with an initial candidate list including all residues within 10 Å of both residue His220 and Lys227 in the compact prestin structure (PDB:7SUN), which are the residues equivalent to the pendrin EC site residues Gln230 and Lys237. A pairwise distance between all Cl^-^ ions and any heavy atom from the candidate residues was calculated, with any distance below 5 Å considered indicative of binding. For each of these residues we calculated the coordination number as the ratio of the time that Cl^-^ ions were bound to this residue to the time they were bound to any of these residues.

The binding site distance *d*_EC,IC_ between the extracellular and intracellular binding sites was calculated as the distance between the C*_α_* atoms of the two closest residues in each site, namely His220–Pro137 in gerbil prestin and Gln230–Pro141 in mouse pendrin.

For all distance calculations, the periodic boundary condition-aware fast distance functions in MDAnalysis ^85^ were used.

### Densities

For all density calculations, trajectories were first structurally superimposed on a common reference frame. For linear densities of lipids and water, the same superposition as for the area analysis was employed, namely superposition on the center of mass of the bilayer, so that area profiles *A*(*z*) and densities *ρ*(*z*) could be directly compared. The `LinearDensity` tool in MDAnalysis ^85^ was used with a histogram bin size of 0.25 Å. For 3D densities of water and Cl^−^ ions, trajectories were superimposed on the transmembrane domain of the protomer of interest and densities collected with the `DensityAnalysis` tool in MDAnalysis ^85^ with a cubic voxel size of 1 Å.

### Molecular visualization

Molecular graphics images were created with UCSF ChimeraX (https://www.cgl.ucsf.edu/ chimerax/) ^39^ and VMD (https://www.ks.uiuc.edu/Research/vmd/)^90^.

## Data and code availability

The structure of prestin with bound thiocyanate was deposited in the Protein Databank with the following accession codes: PDB:9YAY, extended PDB ID pdb_00009YAY, and EMD-72742. The simulation data are available as subsampled trajectories together with input files in an OSF repository with DOI 10.17605/OSF.IO/BXFSA. The Protein_Area code is available from the GitHub repository https://github.com/Becksteinlab/Protein_Area. The analysis code is available from the GitHub repository https://github.com/Becksteinlab/Prestin_Analysis.

## Acknowledgments

The authors wish to thank Dr. Fred Sigworth for his advice and helpful comments. We would like to thank Dr. Marc Llaguno, Dr. Kaifeng Zhou, Dr. Kangkang Song, and Dr. Jianfeng Lin for their assistance with access to the TEM infrastructure and for managing Yale’s electron microscopy facilities. We also thank Dr. Wei Mi for his support in data processing.

Anton 2 computer time was provided by the Pittsburgh Supercomputing Center (PSC) under allocation MCB180088P through grant R01GM116961 from the National Institutes of Health; the Anton 2 machine at PSC was generously made available by D.E. Shaw Research. The authors acknowledge Research Computing at Arizona State University and the Yale Center for Research Computing for providing high performance computing resources that have contributed to the research results reported within this paper. This work utilized the electron microscopy facilities at the Yale School of Medicine.

## Author contributions

C.Z. and R.M. performed MD simulations. C.Z. conceived and wrote the Protein_Area software and analyzed all MD simulations. R.M. performed cryo-EM structure determination and analysis. J.Y. and J.P.B performed electrophysiology. J.Y., J.P.B. and J.S.S. analyzed electrophysiology data. J.S.S., D.S.N., and O.B. conceived the study. C.Z., D.S.N, J.S.S. and O.B. wrote the manuscript with input from all authors.

