## Supplementary Information for "Conformational changes underlying electromechanical transduction in prestin resemble a transport transition in pendrin"

### Signed rotation angles from optimal superposition

We sought to quantify the rotation of the core domain of prestin or pendrin relative to a reference structure, which we defined as the pendrin structure in the inward-facing conformation with PDB ID 7WK1 [1]. This approach relies on the calculation of the rotation matrix after optimal superposition using the QCP algorithm [2], from which the rotation angle $\phi'_i$ and the rotation axis $\hat{\mathbf{n}}_i$ are obtained. The same physical rotation can be equivalently described by an angle $-\phi'_i$ and axis $-\hat{\mathbf{n}}_i$. The optimal superposition algorithm does not return rotations in a fixed reference frame, in other words, the returned rotation axis is not fixed and thus the sign of the rotation angle is not well-defined. In order to use a *signed* rotation angle as an order parameter, one must choose a reference direction for the axis. For small rotations near 0, *i.e.*, the nearly degenerate case when a conformation is close to the reference structure, the rotation axis is poorly defined and its direction is ambiguous and cannot be simply resolved by comparison with a static reference direction. Okazaki et al. [3] sidestepped this issue by always choosing the absolute value of the rotation angle, thus introducing ambiguity in the meaning of $\phi$ while preserving the magnitude as measure of the size of the rotation. Here we introduce a sign correction procedure that is robust to the sign ambiguity in the rotation axis and allows us to use the rotation angle as a meaningful order parameter even near 0.

We chose as the reference direction $\hat{\mathbf{m}}$ the axis of rotation for the rotation of the core domain of OF pendrin (PDB:7WLE) onto the core domain of IF pendrin (PDB:7WK1) that produces a negative rotation angle. For each trajectory frame, we compute the rotation angle $\phi'$ and axis $\hat{\mathbf{n}}$ from the rigid-body rotation matrix R; alternatively and equivalently, this rotation can be described by the unit quaternion $q = (w, \mathbf{v}) = (\cos\frac{\phi'}{2}, \hat{\mathbf{n}}\sin\frac{\phi'}{2})$ (or $-q$). We then project this rotation on a rotation around the reference axis $\hat{\mathbf{m}}$ to obtain the projected rotation angle $\theta$ by computing the unit quaternion $p(\theta) = (\cos\frac{\theta}{2}, \hat{\mathbf{m}}\sin\frac{\theta}{2})$ that describes the closest rotation to $q$ while using axis $\hat{\mathbf{m}}$. To do so, we seek to determine $\theta$ such that it minimizes the relative rotation

$$d(\theta) = qp(\theta)^{-1} \tag{S1}$$

between $q$ and $p(\theta)$. (Note that $q$ transforms a 3-vector $\mathbf{x} \xrightarrow{q} \mathbf{x}'$ as $\mathbf{x}' = q\mathbf{x}q^{-1}$; if using the rotation $p$, a relative rotation $d$ must be added ($\mathbf{x} \xrightarrow{p} \mathbf{x}'' \xrightarrow{d} \mathbf{x}'$ with $\mathbf{x}' = dp\mathbf{x}(dp)^{-1}$) to arrive at the same total rotation $q = dp$ and thus equation (S1) follows.) Minimizing the relative rotation described by $d = (\cos\frac{\Delta}{2}, \hat{\mathbf{u}}\sin\frac{\Delta}{2})$ means to minimize the relative rotation angle $\Delta$ between $q$ and $p(\theta)$ or to maximize the scalar or real part of $d$, $f(\theta) = \Re(d(\theta)) = \cos\frac{\Delta}{2}$ (for $0 \leq \Delta \leq \pi$, as we need cos to be monotonous and we choose to describe the relative rotation by $d$ with $\Re(d) \geq 0$ instead of $-d$). Carrying out the quaternion multiplication in equation (S1), where $p(\theta)^{-1} = \overline{p(\theta)}$ is the quaternion conjugate because $p(\theta)$ is a unit quaternion, and retaining the real part, we obtain

$$f(\theta) = \Re(d(\theta)) = q \cdot p(\theta) = w\cos\frac{\theta}{2} + \mathbf{v}\cdot\hat{\mathbf{m}}\sin\frac{\theta}{2} \tag{S2}$$

where $q \cdot p(\theta)$ denotes the Euclidean inner product of the two quaternions viewed as 4-vectors. We maximize $f(\theta)$ by solving

$$\frac{df(\theta)}{d\theta} = -w\sin\frac{\theta}{2} + \mathbf{v}\cdot\hat{\mathbf{m}}\cos\frac{\theta}{2} = 0 \tag{S3}$$

for $\theta$ as

$$\tan\frac{\theta}{2} = \frac{\sin\frac{\theta}{2}}{\cos\frac{\theta}{2}} = \frac{\mathbf{v}\cdot\hat{\mathbf{m}}}{w}. \tag{S4}$$

equation (S4) determines $\frac{\theta}{2}$ only up to multiples of $\pi$ and therefore has two solutions, namely the maximum and the minimum of $f(\theta)$. At either stationary point, we can write $\sin\frac{\theta}{2} = \alpha\mathbf{v}\cdot\hat{\mathbf{m}}$ and $\cos\frac{\theta}{2} = \alpha w$ with the proportionality constant $\alpha$. The constant is fixed by the trigonometric identity $\sin^2\frac{\theta}{2} + \cos^2\frac{\theta}{2} = 1$ to $\alpha = \pm 1/\sqrt{w^2 + (\mathbf{v}\cdot\hat{\mathbf{m}})^2}$ and thus equation (S2) evaluates to $f = \alpha\left(w^2 + (\mathbf{v}\cdot\hat{\mathbf{m}})^2\right) = \pm\sqrt{w^2 + (\mathbf{v}\cdot\hat{\mathbf{m}})^2}$. The maximum that we seek is the solution $\theta_{\text{max}}$ for which $\cos\frac{\theta}{2}$ has the same sign as $w$ and $\sin\frac{\theta}{2}$ the same sign as $\mathbf{v}\cdot\hat{\mathbf{m}}$ ($f > 0$); the second solution, $\theta_{\text{max}} \pm 2\pi$, is the minimum ($f < 0$, corresponding to $\Delta > \pi$, which we excluded above). The ratio on the right-hand side of equation (S4) does not retain the two signs separately, so the single-argument arctan function cannot distinguish the two solutions and returns the minimum whenever $w < 0$. The two-argument $\arctan 2$ function keeps both signs and hence selects the maximum; it is also numerically robust for $w = 0$, where the ratio diverges and $\theta = \pm\pi$. We therefore obtain

$$\theta = 2\arctan 2(\mathbf{v}\cdot\hat{\mathbf{m}}, w). \tag{S5}$$

Because $q$ and $-q$ describe the same rotation, flipping the sign of the quaternion also flips the signs of $w$ and $\mathbf{v}$ and hence shifts equation (S5) by $\pm 2\pi$; both values describe the same rotation about $\hat{\mathbf{m}}$ but only one of them lies in the interval $-\pi \leq \theta \leq \pi$. We therefore fix the representative by requiring $w \geq 0$, which restricts the projected rotation angle to $-\pi \leq \theta \leq \pi$.

For each trajectory frame $1 \leq i \leq N$, the projected rotation angle $\theta_i$ was calculated according to equation (S5) and used to resolve the sign ambiguity in $\phi'_i$. Because the rotation magnitude can be estimated robustly whereas its sign remains ambiguous, we defined a corrected signed series as

$$\phi_i = s_i \phi'_i, \tag{S6}$$

where $s_i \in \{-1, +1\}$ represents the sign state at frame $i$. We selected the sign path that minimized the objective function

$$L(\{s_i\}_{1\leq i\leq N}) = \sum_{i=1}^{N} \left(\phi_i(s_i) - \theta_i\right)^2 + \gamma_{\text{smooth}} \sum_{i=2}^{N} \left(\phi_i(s_i) - \phi_{i-1}(s_{i-1})\right)^2 . \tag{S7}$$

The first term enforces agreement between the corrected raw angle and the projected rotation angle, whereas the second term penalizes abrupt frame-to-frame changes and therefore favors temporally smooth sign assignments without large jumps that would imply unphysically fast changes in the conformation. Because each frame has only two possible sign states, the optimization can be written as a two-state path problem and solved exactly by dynamic programming. For each frame and candidate sign state, the cumulative cost was computed from the minimum of the two possible predecessor states plus the local data-fit and regularization terms. The globally optimal sign sequence was then recovered by backtracking from the minimum-cost terminal state and $\phi_i$ (equation (S6)) was recorded as the value of the order parameter (Figure S13).

This formulation preserves the magnitude of the original raw angle series exactly because the correction changes only the sign. We verified numerically that the absolute value of the corrected series matched the absolute value of the raw series at every frame. The smoothness hyperparameter $\gamma_{\text{smooth}}$ was evaluated empirically over a broad range of values $10 \leq \gamma_{\text{smooth}} \leq 10^4$ by measuring the “roughness” of the corrected series, here simply defined as the average of the frame-to-frame squared differences $r_i = (\phi_i - \phi_{i-1})^2$ over the largest 0.01% of the values $r_i$ (Figure S14). The percentile filter increased the signal-to-noise ratio because only the larger jumps in the timeseries were considered. Based on these diagnostics, $\gamma_{\text{smooth}} = 1000$ was adopted for the final production workflow as it was the smallest tested value that suppressed all unrealistic sign jumps.

### Protein area calculation

The Protein_Area algorithm uses 2D Voronoi tessellation on slices of the full MD simulation system to calculate the protein area profile $A(t, z)$ at each trajectory frame. In order to handle periodic boundary conditions in the $X$-$Y$ plane, the simulation box is replicated in the $X$-$Y$ plane so that the 2D Voronoi tessellation is periodic. Therefore, only orthorhombic simulation boxes are supported. The system should be oriented such that the membrane normal is along the $Z$ axis. The user has to pre-process the trajectory to center the simulation system on protein center of mass to maintain a consistent reference frame along the membrane normal; a structural superposition is not necessary (and not recommended unless it is restricted to rotations in the $X$-$Y$ plane). In order to find the dividing surface between protein and surrounding medium, the trajectory must contain all atoms. At the moment, the code automatically selects "protein" atoms according to the MDAnalysis "protein" selection[4], which is based on a dictionary of protein residue names, and classifies all other atoms as non-protein.

For each trajectory frame, the protein area profile $A(t,z)$ is calculated by the following steps:

1. The simulation box is sliced on the $Z$ axis at a regular distance interval (by default 0.5 Å for all-atom simulations).
2. For simulations under periodic boundary conditions, the simulation box is replicated in the $X$-$Y$ plane so that the 2D Voronoi tessellation is periodic.
3. For each 2D slice at location $z$, the atoms are assigned to a Voronoi cell (see Figure S15 for an example).
4. The protein area profile $A(t,z)$ is the sum of the areas of all Voronoi cells that were generated from protein atoms.

The code returns the timeseries of the protein area profile $A(t,z)$. The protein area profile $A(t,z)$ is then averaged over all trajectory frames to obtain the final protein area profile $A(z)$.

The Protein_Area Python package uses MDAnalysis[4] to read the trajectory, the Voronoi tessellation is performed with `scipy.spatial.Voronoi` from the SciPy package https://scipy.org/[5], and the area calculation is performed with `shapely.geometry.Polygon` from the shapely package https://github.com/shapely/shapely. The code is available at https://github.com/Becksteinlab/Protein_Area under the GPL-v2 open source license. For this work, we used version 0.1.0.

### Detailed structural characterization of prestin and pendrin

The structural model of expanded prestin obtained from the clustering of the MD simulations (main text Figure 2a) resembles the experimental structure of IF pendrin (PDB:7WK1) more than the compact prestin conformation (PDB:7SUN), as shown by a transmembrane domain (TMD) $C_\alpha$ RMSD of 2.3 Å relative to PDB:7WK1 compared to 3.1 Å relative to PDB:7SUN (Table S2). This similarity is even more evident in the RMSD of the core domain after superposition on the gate domain of 2.8 Å for PDB:7WK1 compared to 5.1 Å for PDB:7SUN (Table S3). The model of the compact-2 state (main text Figure 2b) is structurally intermediate between the compact prestin structure (PDB:7SUN) with a TMD $C_\alpha$ RMSD of 2.7 Å and the pendrin OF structure (PDB:7WLE) with 4.0 Å (Table S2), consistent with the corresponding core domain RMSDs of 4.5 Å for PDB:7SUN and 6.5 Å for PDB:7WLE (Table S3). The gate domain generally superimposes well between all structures and the computational models, *e.g.*, with an RMSD of 1.9 Å between expanded prestin and IF pendrin (PDB:7WK1) or 1.8 Å between compact-2 prestin and OF pendrin (PDB:7WLE; Table S3). The gate domains of the experimental structures superimpose even closer with the largest RMSD of 1.7 Å between compact prestin (PDB:7SUN) and OF pendrin (PDB:7WLE; Table S3). Thus, the expanded prestin conformation resembles the IF conformation of pendrin and the compact-2 prestin conformation appears to be an intermediate conformation between the compact prestin structure and the OF conformation of pendrin. The major conformational difference between the expanded prestin conformation and the IF conformation of pendrin is the larger core-domain shift and larger core rotation in the latter whereas differences in the gate domain are less pronounced.

### Cryo-EM structure characterization

The reconstruction revealed a homodimeric assembly refined with C2 symmetry, in which two identical protomers adopt the characteristic 7+7 inverted repeat fold of the SLC26 transporter family. The cryo-EM density map shows well-resolved transmembrane helices, enabling reliable model building and refinement of the protein structure. We noted poorer resolution at the tips of the extracellular loops, the cytosolic IVS loop (intervening sequence), and parts of the cytosolic C-terminal domain, likely reflecting increased flexibility in these regions. The solved cryo-EM structure has two protomers; each protomer consists of an intracellular N terminus, 14 transmembrane helices (TM1–TM14), and an intracellular C terminus that includes the STAS domain (main text Figure 5a). The transmembrane domain consists of a core domain and a gate domain, forming the canonical architecture observed in previously reported prestin structures, including PDB:7SUN[6] and PDB:7LGU[7]. Eight transmembrane helices (TMs 1–4 and TMs 8–11) constitute the core domain, whereas the remaining helices (TMs 5–7 and TMs 12–14) form the gate domain. Overall, this structure in the presence of $SCN^-$ (PDB:9YAY) most closely resembled our previous structure of prestin in the presence of $Cl^-$ (PDB:7SUN)[6]. In turn, both prestin structures (PDB:9YAY and PDB:7SUN) most closely resemble the intermediate structure of SLC26A9 (PDB:6RTF)[8], with RMSD values of 1.042 and 0.986 Å, respectively.

Inspection of the density map revealed a well-defined density within the internal anion binding pocket, corresponding to the canonical binding site previously determined for chloride. We identified the density as an $SCN^-$ anion within 4 Å of residues Gln97, Phe137, Val221, Ser396, Leu397, Ser398, and Val444 that stabilize the ligand within the binding cavity (Figure S7, panel **e**). These residues have previously been identified in chloride binding[7,9], confirming that this region serves as the conserved internal anion interaction site in prestin.

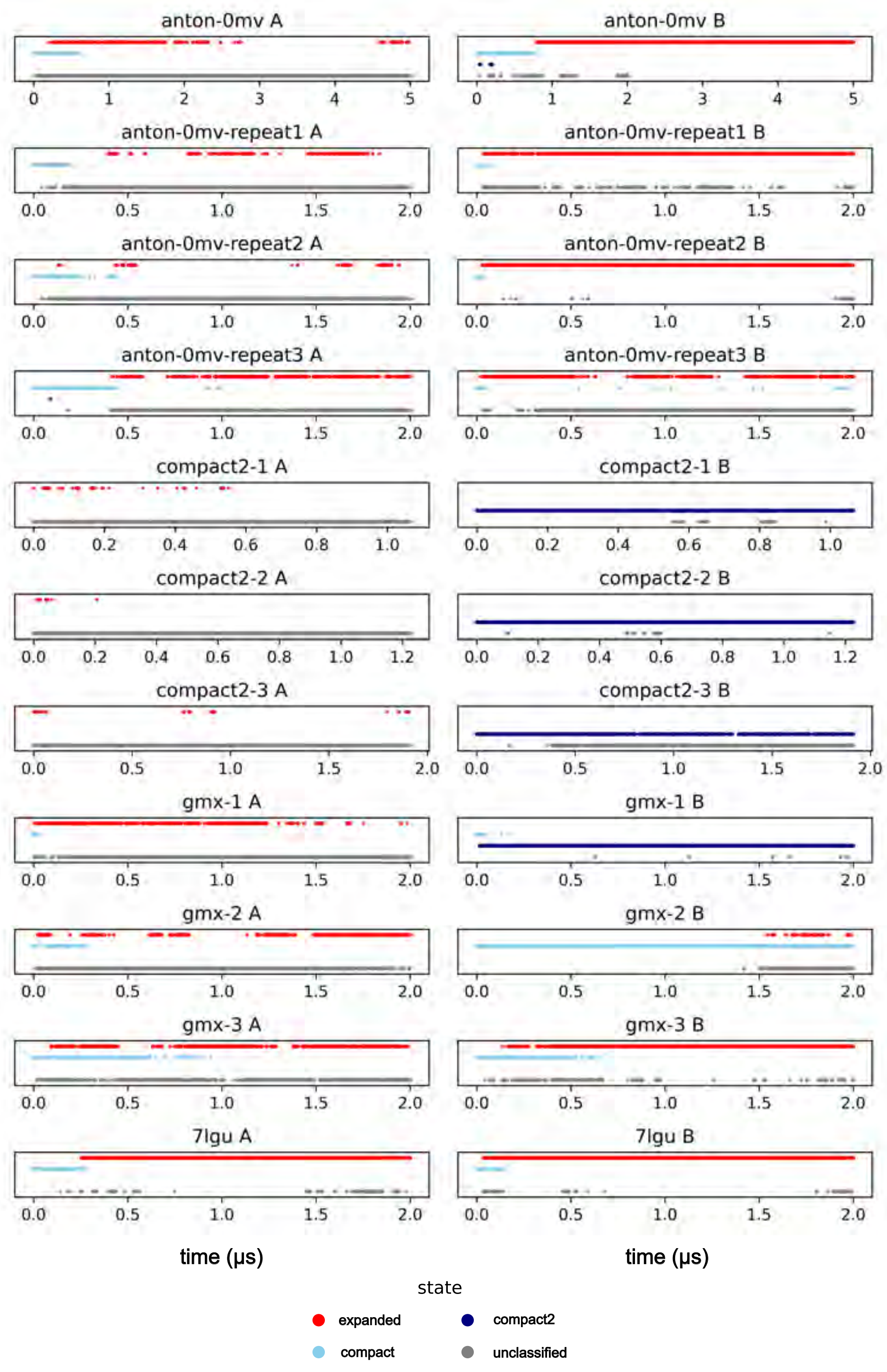


**Figure S1. Classification of prestin conformational states for protomers A and B in MD simulations.** The three structural states are defined as expanded (red), compact (light blue), and compact-2 (dark blue); compact and compact-2 together constitute the contracted macrostate C, whereas expanded constitutes E. Conformations that could not be classified into any of the three states are labeled as “unclassified” (gray). The time axis is adjusted to the different simulation lengths.

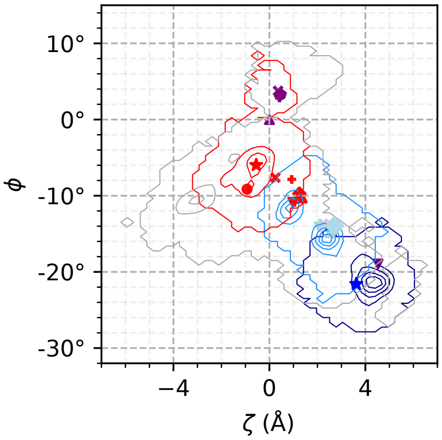


| state | color | count | fraction |
|---|---|---|---|
| expanded | red | 70349 | 24% |
| compact | light blue | 33893 | 11% |
| compact-2 | dark blue | 50044 | 17% |
| unclassified | gray | 141462 | 48% |
| total | | 295748 | 100% |

**Figure S2. State distribution in prestin MD simulations.** For each trajectory frame in all simulations of gerbil prestin, a state label was assigned to protomers A and B using the GROMOS classifier as described in Methods (red: expanded, light blue: compact, dark blue: compact-2). Some conformations are structurally too distant from the reference structures and are not classified into any of the three states and are labeled as “unclassified” (gray). See Figure 2 in the main text for the definition of indicated structures. The outermost contour lines represent all data points with the label.

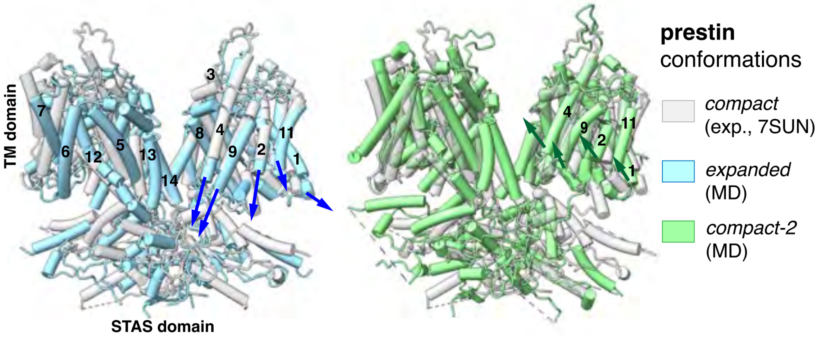


**Figure S3. Prestin domain movement.** (Left) The transition from the compact cryo-EM structure (PDB:7SUN, white) to the expanded conformation (light blue) involves rotation and downward movement of the core domain relative to the gate domain, towards the intracellular side. TM helices 4, 9, 2, 11, and 1 displace lipids in the inner leaflet. (Right) The transition from the compact cryo-EM structure (PDB:7SUN, white) to the compact-2 (green) conformation involves rotation and upward movement of the core domain relative to the gate domain, towards the extracellular side, without noticeable displacement of lipids. Structures were superimposed on the gate domain of the right protomer. Images were produced with ChimeraX [10].

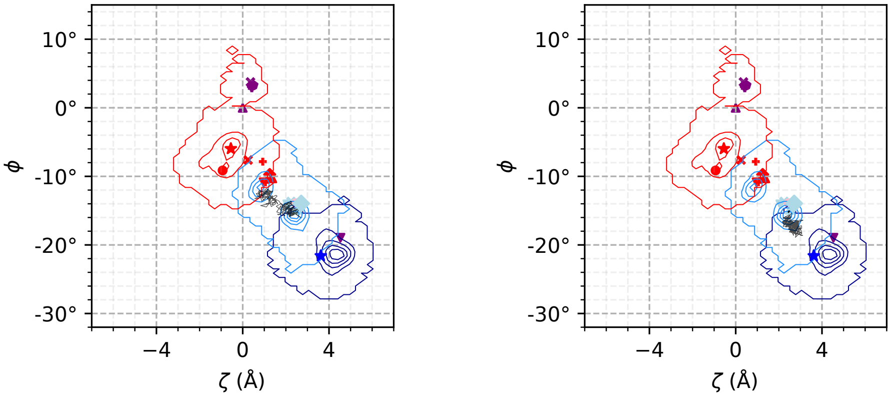


**(a)** Protomer A **(b)** Protomer B

**Figure S4. Projection of the initial time steps of the prestin-Anton-0mV simulation of gerbil prestin onto the $\zeta$-$\phi$ order parameter space.** The simulation started from the compact cryo-EM structure (PDB:7SUN, light-blue diamond ◆) with initial energy minimization and equilibration steps (not shown). The trajectory is shown as a black line for the first 240 ps. Steps are plotted every 1.2 ps. The original Anton2 trajectory was configured with a write-out interval of 240 ps. In order to obtain the higher frequency sampling to resolve any conformational changes in the initial phase of the simulation, we made use of Anton2's exact restart feature that generates identical trajectories from identical initial conditions. We relaunched the prestin-Anton-0mV simulation with a write-out interval of 1.2 ps and total run time of 10 ns. Prestin conformational states in this space are indicated by contour lines (compact: light blue, expanded: red, compact-2: dark blue). See Figure 2 in the main text for the definition of the other markers and the full 5-μs trajectory of protomer B.

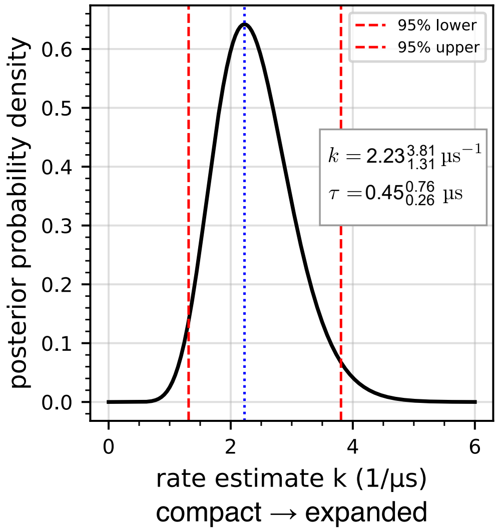


**Figure S5. Prestin conformational change rate estimate from Bayesian inference.** The posterior distribution of the transition rate $k$ from the compact to the expanded state is shown together with the 95% confidence interval. No reverse transition from expanded to either contracted state was observed in any simulation, and hence no reverse rate was estimated. See Table S1 for the gerbil prestin data used for the Bayesian inference.

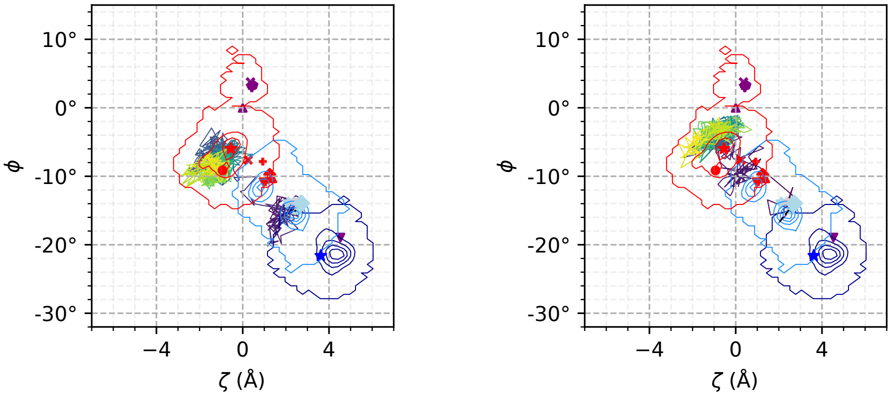


**(a)** Protomer A

**(b)** Protomer B

**Figure S6. Projection of the prestin-7lgu simulation of human prestin onto the $\zeta$-$\phi$ order parameter space.** The simulation started from the compact cryo-EM structure (PDB:7LGU, light-blue upright triangle ▲) with initial energy minimization and equilibration steps, which are indicated as a black dashed line (– – – ). Starting with the equilibrated structure, the trajectory is colored from dark blue (0 µs) to yellow (2 µs) according to simulation time. Steps are plotted every 2.4 ns. Prestin conformational states in this space are indicated by contour lines (compact: light blue, compact-2: dark blue, expanded: red). See Figure 2 in the main text for the definition of the other markers.

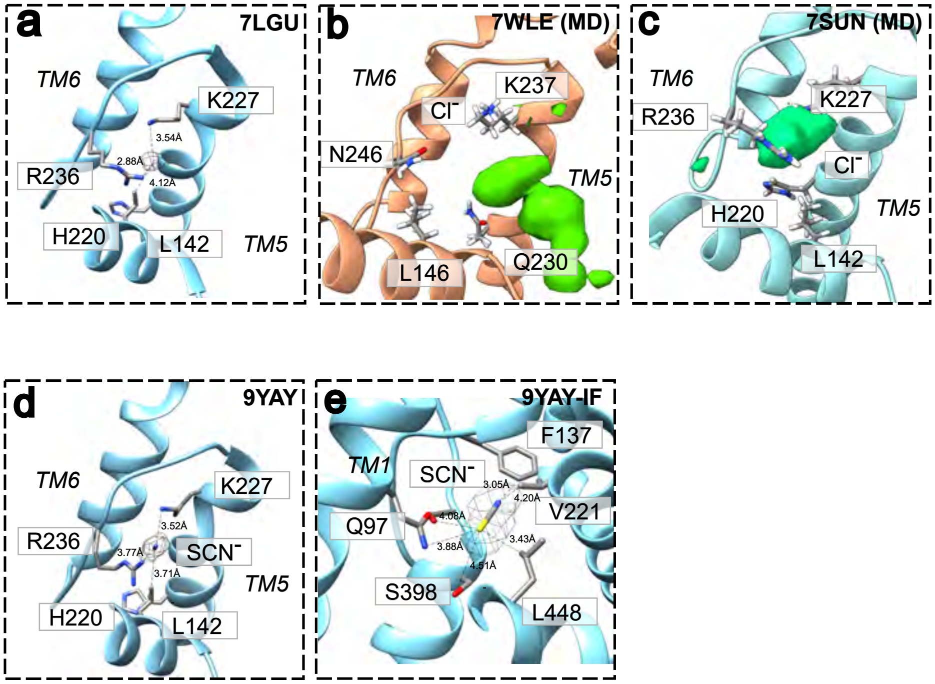


**Figure S7. Anion binding site in prestin and pendrin. a** Isolated density in the cryo-EM map (EMD-23329) of the human 2.3-Å resolution structure (PDB:7LGU) of prestin in the *extracellular anion site* with the density contoured at a Chimera contour level of 0.1. **b** $Cl^-$ density (green) in the *extracellular site* from the pendrin-outward simulation, which was initialized from the OF pendrin structure (PDB:7WLE) in the presence of 150 mM NaCl. The density is contoured at 1.5 M. **c** $Cl^-$ density (green, contoured at 1.5 M) in the *extracellular site* from the prestin-Anton-0mV simulation in the presence of 150 mM NaCl. **d** $SCN^-$ in the *extracellular site* of prestin from the density in the cryo-EM map of gerbil prestin in the presence of thiocyanate (PDB:9YAY), with the ligand density contoured at 0.1. **e** $SCN^-$ in the *intracellular site* in the thiocyanate structure (PDB:9YAY, density contour at 0.1). Images were produced with ChimeraX[10].

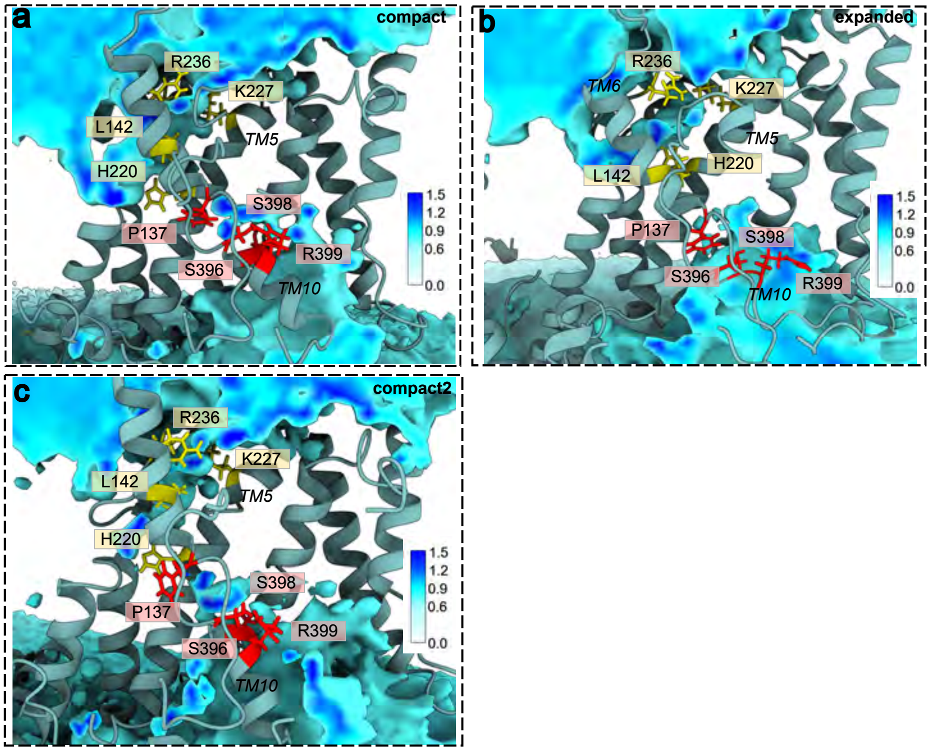


**Figure S8. Water density from MD simulations of prestin in the compact, expanded, and compact-2 conformations.** The compact simulation was initialized from the compact prestin structure (PDB:7SUN). **a** Water density of prestin compact. **b** Water density of prestin expanded. **c** Water density of prestin compact-2. Some helices were removed for clarity. Residues belonging to the canonical intracellular binding site (residues Pro137, Ser396, Ser398, Arg399) are labeled in red; residues in the extracellular binding site (residues L142, His220, Arg236, Lys227) are labeled in yellow. Images were produced with ChimeraX[10].

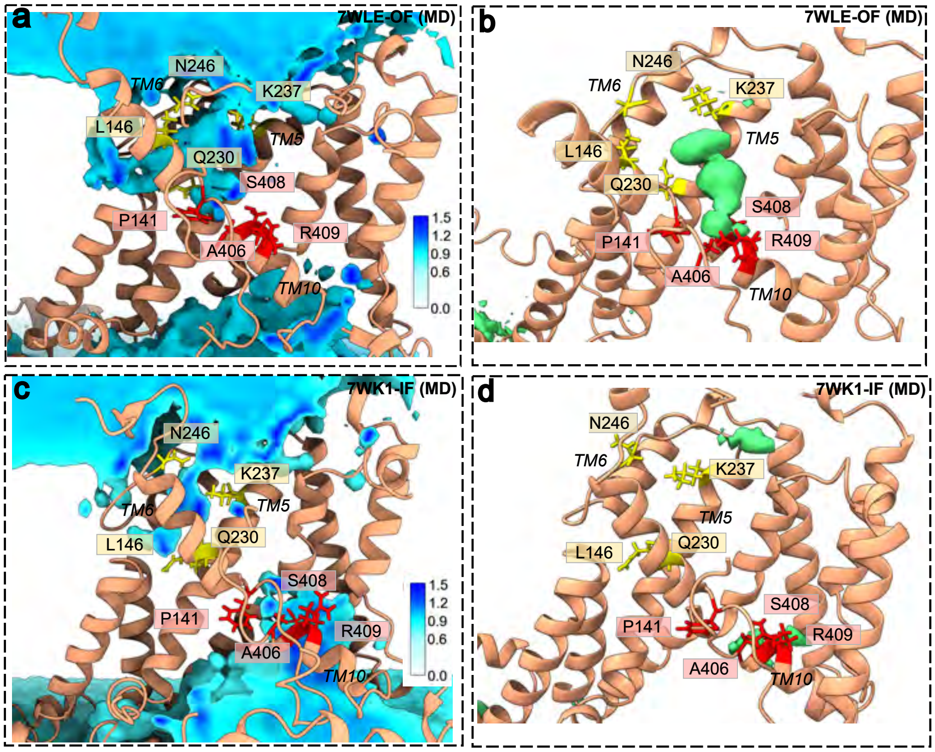


**Figure S9. Water and Cl⁻ density from pendrin MD simulations.** The pendrin-outward simulation was initialized from the OF pendrin structure (PDB:7WLE) while the pendrin-inward simulation was initialized from the IF pendrin structure (PDB:7WK1). **a** Water density of pendrin-outward. **b** Cl⁻ density of pendrin-outward. **c** Water density of pendrin-inward. **d** Cl⁻ density of pendrin-inward. The Cl⁻ density is contoured at 1.5 M. The water densities are scaled relative to the bulk water molarity (55.5 M). Some helices were removed for clarity. Residues belonging to the canonical intracellular binding site (residues Pro141, Ala406, Ser408, Arg409) are labeled in red; residues in the extracellular binding site (residues L146, Gln230, Asn246, Lys237) are labeled in yellow. Images were produced with ChimeraX[10].

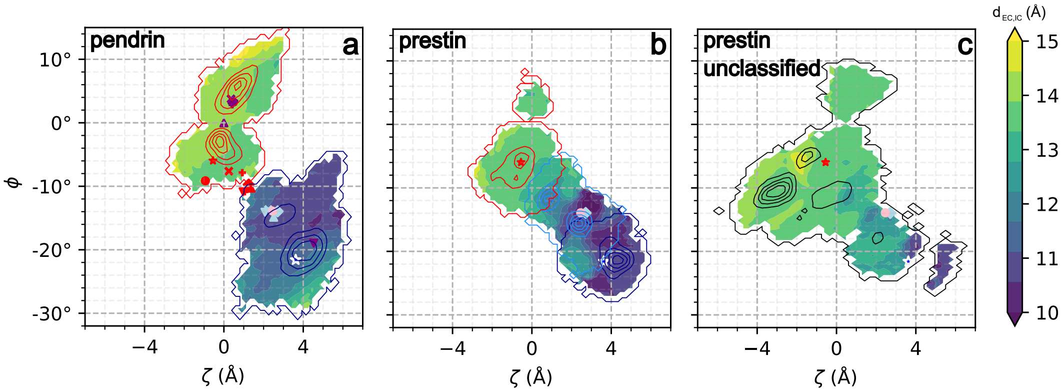


**Figure S10. Anion binding site distance $d_{\mathrm{EC,IC}}$ (in Å) projected onto the $\zeta$-$\phi$ order parameter space. a** Pendrin, all trajectory data (density isocontours for OF simulations in red and for IF simulations in blue; experimental structures are indicated by markers: ▲: pendrin IF conformation (PDB:7WK1), ▼: pendrin OF conformation (PDB:7WLE)) **b** Prestin, only classified trajectory frames (compact (light blue), compact-2 (dark blue), expanded (red); experimental structures ♦: compact (PDB:7SUN), ●: compact (SCN⁻, PDB:9YAY); models from MD: ★: compact-2, ★: expanded) **c** Prestin, only unclassified trajectory frames (black contours). See Figure 2 in the main text for the definition of the other markers.

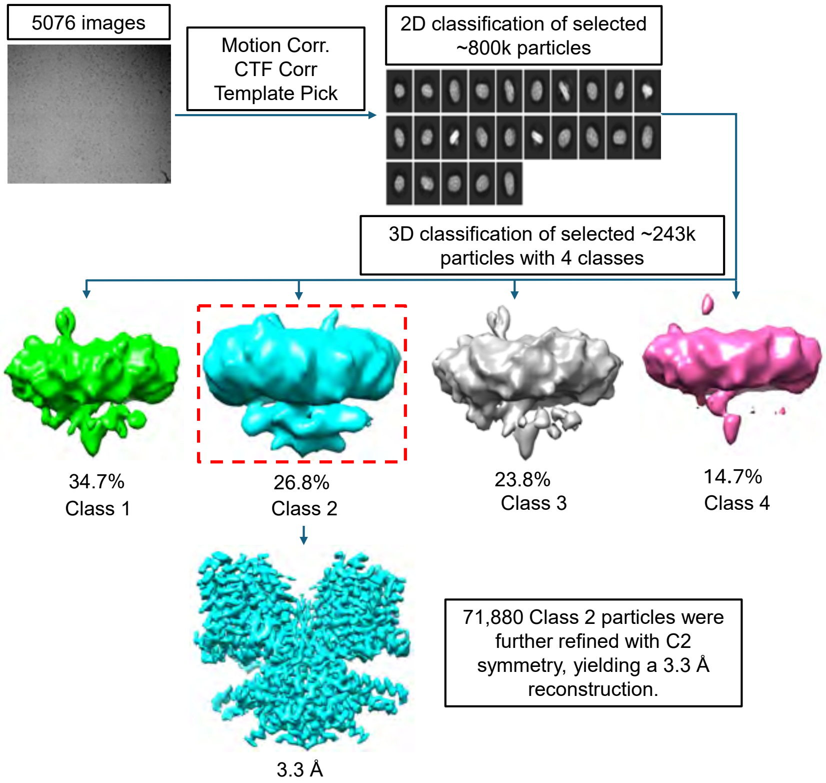


**Figure S11. Cryo-EM data processing workflow for the thiocyanate structure of gerbil prestin.** Data were processed with CryoSPARC and deposited in the Electron Microscopy Data Bank (EMDB) as EMD-72742 and the structural model was deposited in the Protein Data Bank with the accession code pdb_00009YAY.

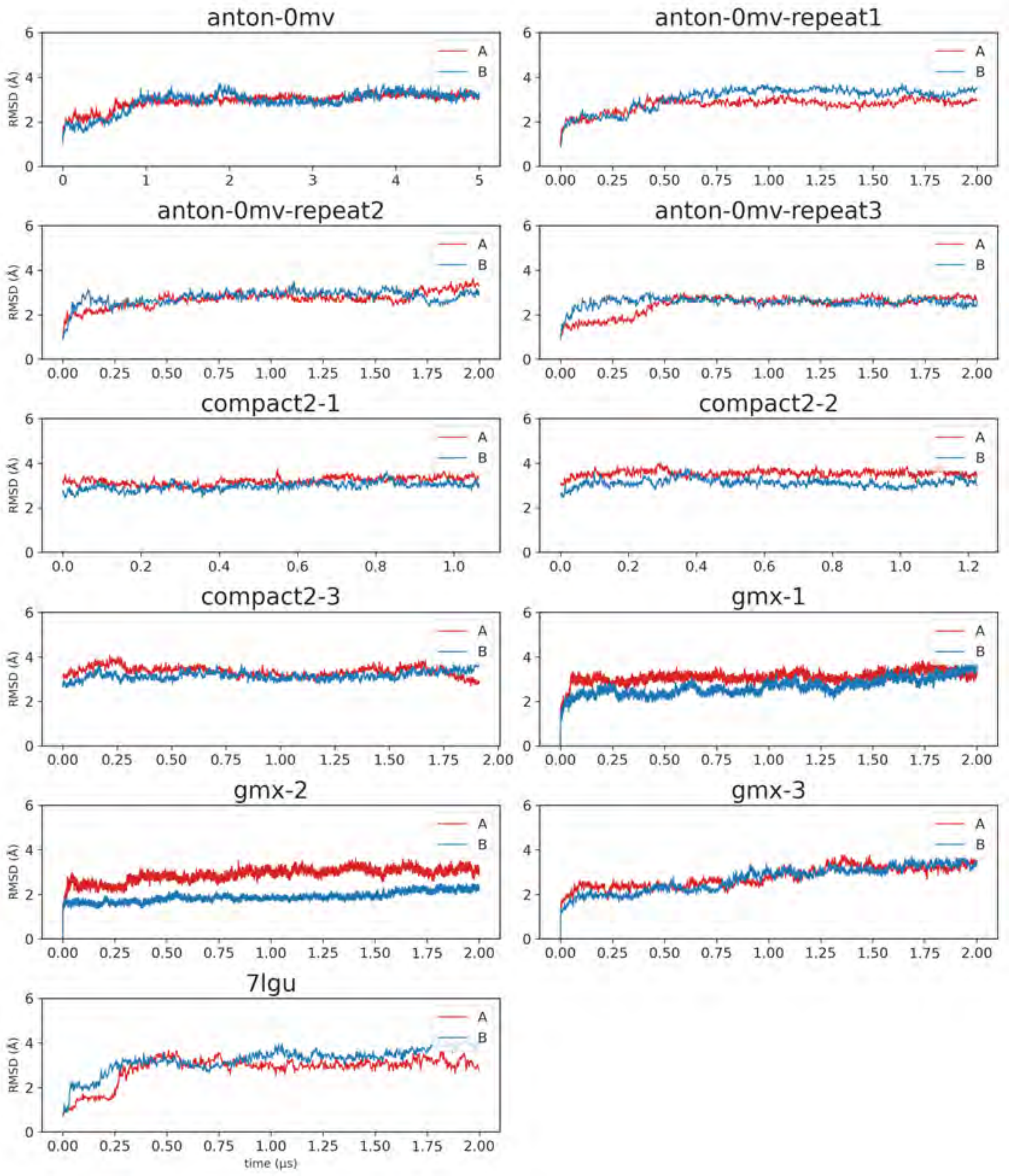


**Figure S12. Transmembrane domain $C_\alpha$ RMSD of prestin simulations.** For gerbil prestin simulations, the $C_\alpha$ RMSD of the transmembrane domain was calculated for each protomer after the protein was aligned to the transmembrane domain (residues 76–155 and 167–504) of gerbil prestin (PDB:7SUN) separately on protomer A and protomer B. The $C_\alpha$ RMSD of the prestin-7lgu simulation was calculated with the same alignment described above to the experimental structure (PDB:7LGU).

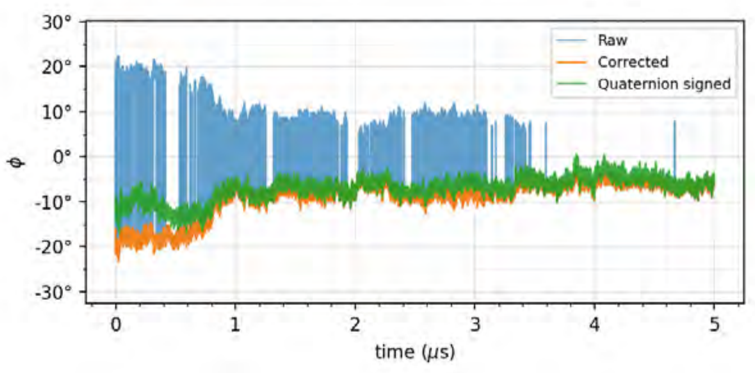


**(a)** Prestin (PDB:7SUN, simulation prestin-Anton-0mv, protomer A)

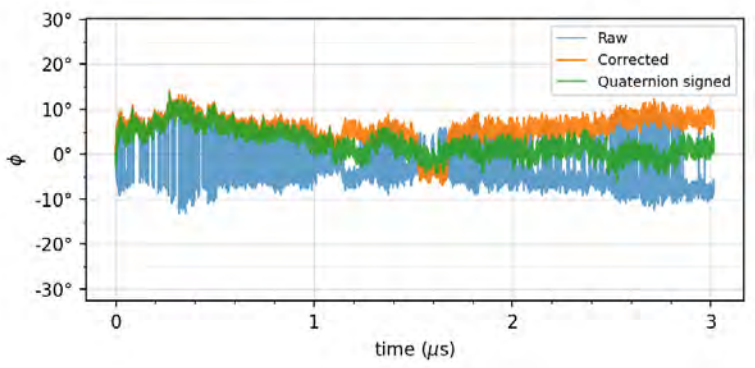


**(b)** Pendrin (PDB:7WK1, simulation pendrin-inward, protomer B)

**Figure S13. Signed rotation angle correction.** The rotation angle time series $\{\phi_i'\}_{1\le i\le N}$ that was directly obtained from the optimal superposition rotation matrix (blue, "Raw") shows random sign changes. The projected angle time series $\{\theta_i\}_{1\le i\le N}$ (green, "Quaternion signed") is obtained by projecting the original rotation onto the fixed reference axis (defined by the rotation of the core domain of OF pendrin (PDB:7WLE) onto the core domain of the reference structure, IF pendrin (PDB:7WK1)). The sign-corrected timeseries $\{\phi_i\}_{1\le i\le N}$ was obtained from $\phi_i'$ and $\theta_i$ by minimizing the objective function equation S7 with a dynamic programming algorithm. It has the same magnitude as the original timeseries but generally follows the projected angle without major unphysical jumps (orange, "Corrected raw (DP)"). **a** Simulation of gerbil prestin (PDB:7SUN), showing a transition from the compact to the expanded state, which is closer to the reference structure; rotation angles are generally large and negative and in principle assignment is not ambiguous. **b** Simulation of mouse pendrin (PDB:7WK1), showing equilibrium fluctuations around the IF reference structure. The overall conformation does not change, with the core domain essentially remaining in place. Therefore, the magnitude of rotations is small and the direction of the rotation depends sensitively on local atomic fluctuations. In some instances the projected angle is close to zero, indicating that the original rotation axis is perpendicular to the reference axis. In this case, the regularization term is important to avoid unrealistic jumps in $\phi_i$.

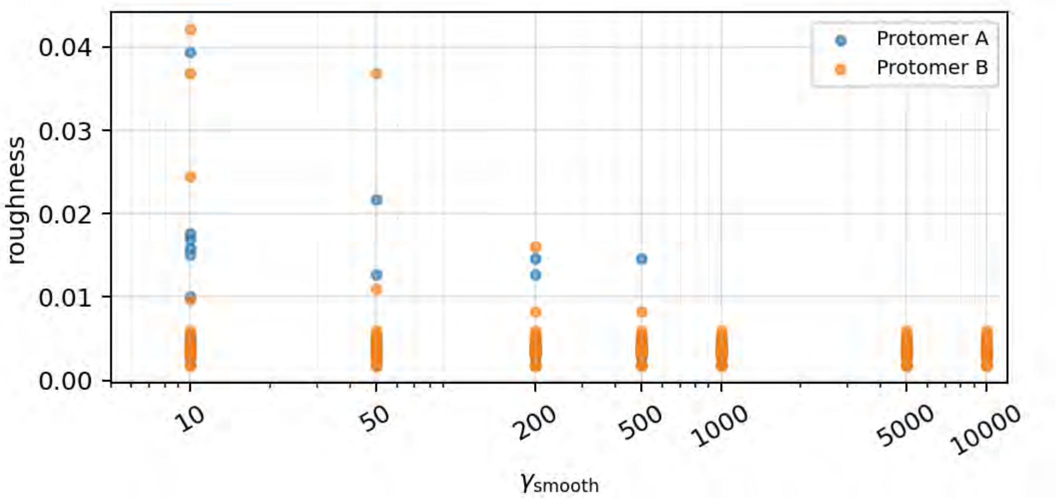


**Figure S14. Selection of the smoothness hyperparameter for the signed rotation angle correction.** The "roughness" $r_i = (\phi_i - \phi_{i-1})^2$ for each trajectory frame $i$ was averaged over the largest 0.01% of $r_i$ values in each timeseries. The smoothness hyperparameter $\gamma_{\mathrm{smooth}}$ was varied from $10$ to $10^4$ and the roughness was computed for each protomer (A, B) in each simulation (Table 1 in the main text). The choice of $\gamma_{\mathrm{smooth}} = 1000$ was the smallest value that left the distribution of roughness values unchanged and suppressed all sign jumps greater than 0.01.

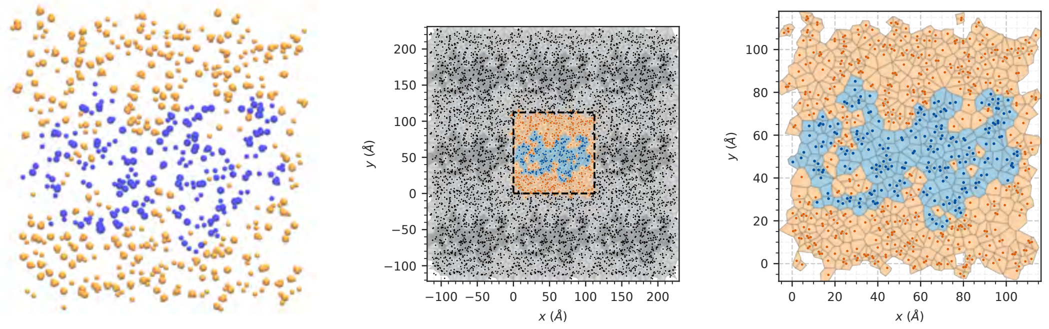


**(a)** atom positions **(b)** periodic Voronoi tessellation **(c)** unit cell Voronoi tessellation

**Figure S15. 2D Voronoi tessellation for membrane protein area profile calculation.** Example of the periodic Voronoi tessellation in one layer of the prestin simulation system. Voronoi cell centers associated with protein atoms are indicated by blue dots and are shaded blue. The medium (water, lipids, ions, etc.) is indicated by orange dots and shaded light orange. The protein area is the sum of the areas of all blue cells. Axes in the $X$-$Y$ plane are in Å. **a** Atom positions (Voronoi centers) in a $\Delta z = 0.5$ Å slice (Image generated with VMD [11]). **b** Voronoi tessellation generated from the atom positions with periodic images included; the primary unit cell is indicated by the black dashed rectangle and cells outside the primary unit cell are shaded gray with black centers. **c** Periodic Voronoi tessellation of the unit cell as ultimately used for the area profile calculation. Note that cells near the border are properly bounded by the periodic images and each center is only counted once.

**Table S1. Waiting times for the transition from the compact to the expanded state.** The waiting time is the time that it took for a protomer, which was initially in the compact state, to first visit the expanded state. If no transition was observed, ∞ is shown and the Bayesian estimation uses the total simulation time (in parentheses) as the waiting time [12]. The prestin simulations that were initialized from prestin-gmx-1 (prestin-compact-{1,2,3}) were excluded because the initial states in both protomers did not match the initial state of the compact → expanded transition. For the rate estimate (Figure S5), only simulations with gerbil prestin were used while prestin-7lgu (human prestin) was excluded.

| simulation ID | protomer A (ns) | protomer B (ns) |
|---|---|---|
| prestin-Anton-0mV | 200.64 | 799.68 |
| prestin-Anton-repeat-1 | 398.16 | 36.0 |
| prestin-Anton-repeat-2 | 129.84 | 28.56 |
| prestin-Anton-repeat-3 | 417.6 | 23.52 |
| prestin-gmx-1 | 8.6 | ∞ (2000) |
| prestin-gmx-2 | 21.5 | 1540.9 |
| prestin-gmx-3 | 93.3 | 137.3 |
| prestin-7lgu | 253.68 | 39.96 |

**Table S2. Global structural similarity between prestin conformations and pendrin structures.** Transmembrane domain (upper triangle) $C_\alpha$RMSD (in Å) for prestin simulations were calculated after optimal structural superposition on the transmembrane domain of a single protomer.

| TM | 7WK1 | expanded | 7SUN | compact-2 | 7WLE |
|---|---|---|---|---|---|
| 7WK1 | | 2.30 | 2.92 | 4.42 | 4.26 |
| expanded | | | 3.10 | 4.28 | 4.32 |
| 7SUN | | | | 2.65 | 2.58 |
| compact-2 | | | | | 3.96 |
| 7WLE | | | | | |

**Table S3. Local structural similarity between prestin conformations and pendrin structures.** Core (upper triangle) and gate (lower triangle) $C_\alpha$ RMSD (in Å) for prestin simulations were calculated after optimal structural superposition on the gate domain of a single protomer.

| core / gate | 7WK1 | expanded | 7SUN | compact-2 | 7WLE |
|---|---|---|---|---|---|
| 7WK1 | | 2.80 | 5.08 | 8.37 | 7.22 |
| expanded | 1.93 | | 5.12 | 8.22 | 7.18 |
| 7SUN | 1.31 | 2.14 | | 4.49 | 5.13 |
| compact-2 | 1.92 | 2.51 | 1.88 | | 6.48 |
| 7WLE | 0.85 | 2.10 | 1.73 | 1.78 | |

**Table S4. Mutations affecting NLC.** Mutations in bold face change $z$ significantly. $P$ values are for Student's $t$-test comparing the mutant to the WT. The WT reference value was measured independently in each study. Oliver et al. [13] did not report explicit $z$ or $P$ values for their mutations.

| mutation | $z$ | $P$ | reference |
|---|---|---|---|
| WT[1] | $0.69 \pm 0.01$ | | this work |
| **K75Q** | $0.64 \pm 0.02$ | $< 0.05$ | this work |
| **K77Q** | $0.61 \pm 0.02$ | $< 0.0001$ | this work |
| **E404Q** | $0.59 \pm 0.02$ | $< 0.0001$ | this work |
| WT | $0.73 \pm 0.02$ | | Bai et al. [14] |
| **R130Q** | $0.67 \pm 0.02$ | $< 0.05$ | Bai et al. [14] |
| **E207Q** | $0.68 \pm 0.02$ | $< 0.05$ | Bai et al. [14] |
| **R211Q** | $0.54 \pm 0.01$ | $< 10^{-8}$ | Bai et al. [14] |
| **K255Q** | $0.65 \pm 0.02$ | $< 0.05$ | Bai et al. [14] |
| **K276Q** | $0.63 \pm 0.03$ | $< 0.001$ | Bai et al. [14] |
| **E280Q** | $0.62 \pm 0.02$ | $< 0.0001$ | Bai et al. [14] |
| **K359Q** | $0.65 \pm 0.02$ | $< 0.01$ | Bai et al. [14] |
| **K364Q** | $0.6 \pm 0.02$ | $< 0.0001$ | Bai et al. [14] |
| **D370N** | $0.69 \pm 0.01$ | $< 0.05$ | Bai et al. [14] |
| **K449Q** | $0.64 \pm 0.01$ | $< 0.001$ | Bai et al. [14] |
| **D457N** | $0.66 \pm 0.02$ | $< 0.01$ | Bai et al. [14] |
| **R463Q** | $0.63 \pm 0.03$ | $< 0.005$ | Bai et al. [14] |
| WT[1] | $0.70 \pm 0.02$ | | this work |
| E78Q | $0.70 \pm 0.02$ | 0.69 | this work |
| R281Q | $0.67 \pm 0.05$ | 0.46 | this work |
| WT | $0.73 \pm 0.02$ | | Bai et al. [14] |
| R150Q | $0.71 \pm 0.02$ | 0.59 | Bai et al. [14] |
| K227Q | $0.74 \pm 0.02$ | 0.93 | Bai et al. [14] |
| E433Q | $0.75 \pm 0.01$ | 0.56 | Bai et al. [14] |
| D485N | $0.71 \pm 0.03$ | 0.78 | Bai et al. [14] |
| D154N | | | Oliver et al. [13] |
| D155N | | | Oliver et al. [13] |
| E169Q | | | Oliver et al. [13] |
| R197Q | | | Oliver et al. [13] |
| K233Q[2] | | | Oliver et al. [13] |
| K235Q[2] | | | Oliver et al. [13] |
| R236Q[2] | | | Oliver et al. [13] |
| E277Q | | | Oliver et al. [13] |
| K285Q[3] | | | Oliver et al. [13] |
| E284Q[3] | | | Oliver et al. [13] |
| K283Q[3] | | | Oliver et al. [13] |
| D332Q | | | Oliver et al. [13] |
| D342Q | | | Oliver et al. [13] |
| K409Q | | | Oliver et al. [13] |

[1] WT was measured as part of the same set of experiments as the mutants in this section of the table.

[2] Cluster 1 of positively charged residues (C1) in prestin [13]

[3] Cluster 2 of positively charged residues (C2) in prestin [13]

**Table S5. Coordination number in the prestin extracellular binding site.** Probabilities for a $Cl^-$ ion to be within 5 Å of a heavy atom of the residue while the ion was present in the binding site were calculated over all gerbil prestin MD trajectories together with the corresponding standard deviations. Only residues with a coordination number $\geq 0.1$ are shown.

| residue | coordination number |
|---|---|
| D154 | $0.13 \pm 0.06$ |
| S224 | $0.25 \pm 0.19$ |
| K227 | $0.45 \pm 0.21$ |
| Y228 | $0.16 \pm 0.07$ |
| K233 | $0.21 \pm 0.09$ |
| T234 | $0.16 \pm 0.07$ |
| K235 | $0.28 \pm 0.13$ |
| R236 | $0.54 \pm 0.18$ |
| Y237 | $0.14 \pm 0.09$ |
| S246 | $0.16 \pm 0.15$ |

**Table S6. Cryo-EM image processing and model refinement for the thiocyanate structure of gerbil prestin (PDB:9YAY).**

| **Image Processing** | |
|---|---|
| Microscope | Titan Krios |
| Voltage (kV) | 300 |
| Camera | Gatan K3 |
| Magnification | 81,000 |
| Electron exposure($e^-/\text{Å}^2$) | 54 |
| Defocus range ($\mu$m) | $2.0 - 2.5$ |
| Pixel size (Å) | 0.534 |
| Symmetry imposed | C2 |
| Micrographs | 5076 |
| Final particle images | 71,880 |
| Map resolution (Å) | 3.27 |
| Map resolution threshold (FSC) | 0.143 |
| Map sharpening B factor | $-128$ |
| **Model Refinement** | |
| r.m.s deviation bonds (Å) | 0.003 |
| r.m.s deviation angles (°) | 0.68° |
| MolProbity score | 1.64 |
| Clash score | 6 |
| Rotamer outliers(%) | 2 |
| Ramachandran favored (%) | 97 |
| Ramachandran allowed (%) | 3 |
| Ramachandran outliers (%) | 0 |
